# On the taxonomy and nomenclature of the annual species of the *Brachypodium distachyon* grass complex (Gramineae, Pooideae)

**DOI:** 10.1101/2025.05.13.653784

**Authors:** P. Pablo Ferrer-Gallego, Javier Fabado, Pilar Catalán

**Affiliations:** Servicio de Vida Silvestre y Red Natura 2000, Centro para la Investigación y Experimentación Forestal (CIEF), Generalitat Valenciana, Avda. Comarques del País Valencià, 114, 46930 Quart de Poblet, Valencia, Spain; Jardí Botànic de la Universitat de València. C/Quart, 80. 46008 Valencia, Spain; Departamento de Ciencias Agrarias y del Medio Natural. Escuela Politécnica Superior de Huesca. Universidad de Zaragoza. C/ Carretera de Cuarte Km 1. E-22071 Huesca. Spain; Grupo de Bioquímica, Biofísica y Biología Computacional (BIFI, UNIZAR), Unidad Asociada al CSIC

**Keywords:** *Brachypodium stacei*, *Brachypodium hybridum*, *Bromus distachyon*, lectotype, morphology, nomenclature, synonyms, taxonomy, typification

## Abstract

The taxonomy and nomenclature of the type materials potentially attributable to the three recognized annual species of the grass genus *Brachypodium* (*B. distachyon, B. stacei, B. hybridum*) have been investigated through statistical discriminant analysis of twelve quantitative morphological traits in eight herbarium types corresponding to heterotypic synonyms and newly designated types of each species, along with the lectotype and the epitype of *B. distachyon* and the holotypes of *B. stacei* and *B. hybridum*, plus 50 additional representative specimens of the currently recognized taxa, and through a comprehensive nomenclatural review. The possible taxonomic assignments of types to these species and the nomenclatural priority of *Brachypodium distachyon* and ten related names, *B. hybridum*, *B. macrostachyum, B. megastachyum*, *B. stacei*, *Bromus pentastachyos*, *Festuca monostachyos*, *F. rigida*, *Trachynia platystachya*, *Triticum schimperi*, and *T. subtile*, have been discussed. Lectotypes are designated for *Festuca monostachyos*, *F. rigida*, *Triticum schimperi*, and *T. subtile*. Based on our taxonomic and nomenclatural study, the names *Festuca monostachyos* and *Triticum subtile* can be considered conspecific and heterotypic synonyms of the Linnaean name *Bromus distachyos* (≡ *Brachypodium distachyon*). Furthermore, the names *Brachypodium macrostachyum*, *Brachypodium megastachyum*, *Bromus pentastachyos*, *Festuca rigida*, and *Brachypodium distachyon* var. *platystachyon* (≡ *Trachynia platystachya*) are conspecific with *Brachypodium hybridum*, and *Triticum schimperi* is conspecific with *Brachypodium stacei*. Since the names *B. hybridum* and *B. stacei* are in common use, but both are later heterotypic synonyms, respectively, of the names of the species with which they are conspecific, a proposal for the conservation of these two names is necessary. On the other hand, the application of the names *Bromus pauper*, *Brachypodium geniculatum*, *Festuca pseudistachya*, *Bromus paradoxus*, and *Triticum flabellatum* is uncertain, and as such, they are treated as nomen ambiguum. Finally, *Festuca ciliata* and *Triticum asperum* are two illegitimate names, according to Art. 52.1 of the Shenzhen Code, while *Triticum brevisetum* is a new synonym of *Brachypodium retusum*. A lectotype for the name *T. brevisetum* is also selected and designated in this paper.

## Introduction

*Brachypodium* Palisot de Beauvois (1812: 100, 155) (Gramineae, Pooideae) is a genus of temperate grasses distributed worldwide (Smith 1980, Catalán *et al*. 1995, Catalán & Olmstead 2000, Piep 2007, and comprises about 20 species (Schippmann 1991, Catalán *et al*. 2012, 2016a, 2023, Díaz-Pérez *et al*. 2018, POWO 2025).

*Brachypodium* represents a model system for functional genomics and biology of grasses and monocots (Draper *et al*. 2001, Scholthof *et al*. 2018, Hasterok *et al*. 2022). The completion of the *Brachypodium distachyon* (L.) Palisot de Beauvois (1812: 101, 155, 156. [‘*distichyos*’, ‘*distachyum*’]) (≡ *Bromus distachyos* Linnaeus (1756 (10 March): 13); Linnaeus (1756 (2 June): 8); ≡ *Trachynia distachyos* (L.) Link (1827: 41 [‘*distachya*’]) genome (reference line Bd21; The International *Brachypodium* Initiative-IBI 2010), the first pooid genome to be fully sequenced, spawned considerable scientific interest for comparative genomic analyses of this plant and other totally or partially sequenced genomes of grasses (Catalán *et al*. 2014, Vogel 2016, Scholthof *et al*. 2018, Hasterok *et al*. 2022). Today, high quality reference genomes of the other annual *Brachypodium* species (*B. stacei* Catalán *et al*. (2012: 402 [18 of 21]—reference lines ABR114 and Bsta-ECI; *B. hybridum* Catalán *et al*. (2012: 402 [18 of 21]—reference lines ABR113, Bhyb26, Bhyb-ECI) (Gordon *et al*. 2020, Scarlett *et al*. 2023, Mu *et al*. 2023a, 2023b) and of the slender perennial *B. sylvaticum* (Huds.) Palisot de Beauvois (1812: 155, 156, 101, 181, pl. III fig. 11 (≡ *Festuca sylvatica* Hudson (1762: 38)— reference line Bsyl-Ain1) (Lei *et al*. 2024) are available, expanding the use of these model genomes for investigating the recurrent origins and dynamics of allopolyploids and the functional features and the regulation of expression patterns in perennial plants, respectively.

*Brachypodium distachyon* is a grass species native to southern Europe, northern Africa and south-western Asia to Iraq (Schippmann 1991, Catalán *et al*. 2012, 2016a, 2016b, López-Álvarez *et al*. 2017, Minadakis *et al*. 2023). The close relationship of *B. distachyon* with economically important temperate cereals, forage grasses and biofuel crop grasses, combined with many other favourable attributes, such as its very small and compact nuclear genome, simple growth requirements, small size, and annual life cycle, prompted researchers to propose it as a model organism (Draper *et al*. 2001, IBI 2010, Scholthof *et al*. 2018, Hasterok *et al*. 2022). However, pioneering cytogenetic, taxonomic and molecular phylogenetic studies demonstrated that the three cytotypes (2n=10, 20, 30) of the previously considered single species *B. distachyon* did not correspond to an ascendant autopolyploid series of the same species but to three independent species with different chromosome base numbers and ploidy levels (Catalán *et al*. 2012, and references therein). These authors confirmed the taxonomic identity of the first sequenced genome as belonging to *B. distachyon* (2n=10), and described the other taxa as new species *B. stacei* (2n=20) and *B. hybridum* (2n=30).

Therefore, the *Brachypodium distachyon* complex includes three annual species native to the Mediterranean region that are characterized by a short life cycle, ephemeral habit and self-fertility (Catalán *et al*. 2012, 2016a, 2016b, Scholthof *et al*. 2018). This complex consists of two diploids, each with a different chromosome base number: *B*. *distachyon* (*x* = 5, 2*n* = 2*x* = 10) and *B*. *stacei* (*x* = 10, 2*n* = 2*x* = 20), and their derived allotetraploid *B*. *hybridum* (*x* = 5+10, 2*n* = 4*x* = 30) (Catalán *et al*. 2012). Contrary to some misconceptions, the two diploids are not sister taxa but independent lineages, with *B. stacei* being an early-divergent ancestral lineage of *Brachypodium*, and *B. distachyon* being an intermediate, more recently evolving lineage in all phylogenies of the genus reconstructed from different sorts of genomic data (Catalán *et al*. 2012, 2014, Díaz-Pérez *et al*. 2018, Sancho *et al*. 2022). Furthermore, these two diploid lineages display a descending dysploidy trend from an ancestral *Brachypodium* karyotype (x=10, S), present in *B. stacei*, to a more recent reduced karyotype (x=5, D), present in *B. distachyon*, inferred to have occurred through four nested chromosome fusions (Lusinska *et al*. 2019, Sancho *et al*. 2022). Bayesian dated trees based on nuclear transcriptome data have estimated a Mid-Late Miocene age for the *B. stacei* split (10.2 Ma) and a Late Miocene age for that of *B. distachyon* (7.1 Ma) (Sancho *et al*. 2022), while coalescent dated trees based on whole-genome SNPs have confirmed a similar age for the ancestral split and have also uncovered recent Pleistocene ages for the radiations of the two progenitor species and for the multiple origins of the allotetraploid *B. hybridum* (Gordon *et al*. 2020, Mu *et al*. 2023b, Campos *et al*. 2024).

Previous evidence from different molecular sources, like seed protein data (Hammami *et al*. 2011), nuclear SSRs (Giraldo *et al*. 2012; Shiposha *et al*. 2020), DNA barcoding (López-Álvarez *et al*. 2012), isozymes (Jaaska 2014), and nuclear single-copy genes (Diaz-Pérez *et al*. 2018, Sancho *et al*. 2022) confirmed the co-occurrence of progenitor *B*. *distachyon* and *B*. *stacei* markers in *B*. *hybridum*. Exhaustive comparative genomic analyses have further corroborated the high collinearity of the *B. stacei*-type (S) and *B. distachyon*-type (D) subgenomes of *B. hybridum* with its respective progenitor species genomes (Gordon *et al*. 2020). Phylogenomic studies using whole genome sequence data have reassessed the multiple and bidirectional origins of *B. hybridum*, detecting an ancestral lineage, distributed in Spain, showing a maternal *B. distachyon*-type plastome (D- plastotypes), dated 1.4 Ma, and a more recent lineage, distributed mainly in the western Mediterranean area, showing a maternal *B. stacei*-type plastome (S-plastotypes), dated 0.14 Ma (Gordon *et al*. 2020, Scarlett *et al*. 2023). A recent comparative genomic and phylogenomic study of *B. hybridum* samples from its native circum-Mediterranean region has detected a third, independent and also recent origin for this allotetraploid (S-plastotype, 0. 13 Ma) in the eastern Mediterranean area from local ancestors (Mu *et al*. 2023b). *Brachypodium hybridum* therefore represents a fascinating case of multiple origins of the same allotetraploid species from the same progenitor species (*B. stacei*, *B. distachyon*) but different parental genotypes and bidirectional crosses in different geographical environments of the Mediterranean region and at different times.

Although the three species of the *B. distachyon* complex can be differentiated through their unique cytogenetic traits (chromosome numbers, barcoded karyotypes) and molecular, phenotypic and genomic traits (Catalán *et al*. 2012, 2016a, López-Álvarez *et al*. 2012, 2017, Lusinska *et al*. 2019, Diaz-Perez *et al*. 2018, Sancho *et al*. 2022, Gordon *et al*. 2020, Scarlett *et al*. 2023, Mu *et al*. 2023b, Chen *et al*. 2024), their direct identification is not always straight-forward. As with many taxonomically close angiosperm species, wild individuals show overlapping phenotypic variation due to the plasticity of some morphological characters and adaptation to local environments (Catalán *et al*. 2016b).

However, statistical analysis of 15 traits measured in a large representation of individuals from each species, recovered eight discriminant morphometric traits that differentiated among the three species (leaf stomatal guard cell length, pollen grain length, plant height, culm leaf-blade width, panicle length, number of spikelets per panicle, lemma length, and awn length), four between *B. distachyon* vs. *B. stacei* and *B. hybridum* (culm leaf-blade length, total spikelet length, spikelet length —up to the 4^th^ floret—, and caryopsis length), and one between *B. hybridum* vs. *B. distachyon* and *B. stacei* (number of culm nodes) (Catalán *et al*. 2012, 2016b). These authors also examined five qualitative traits and found tree of them able to discriminate *B. stacei vs B. distachyon* and *B. hybridum* (leaf blade shape, leaf blade softness, leaf blade hairiness), and two between *B. distachyon* vs. *B. stacei* and *B. hybridum* (leaf blade color, occasional presence of rhizomes). Morphologically, *B. stacei* separates from the other congeners based on its curled, soft and densely hairy leaf blades and high stature. In addition, *B*. *hybridum* and *B*. *stacei* are, overall, taller and more robust plant species than *B*. *distachyon*, and the allotetraploid (*B*. *hybridum*) shows larger measurements in several traits (leaf stomatal guard cell length, pollen grain length, and number of culm nodes) than either of its diploid parents, a likely consequence of polyploidy and heterosis (Catalán *et al*. 2012, 2016b).

The nomenclature of the species of the *Brachypodium distachyon* complex has been a matter of debate. *B. distachyon* (basionym *Bromus distachyos* L.) was neotypified by Schippmann and Jarvis (1988) with the specimen LINN no. 93.48. Schippmann (1991) recorded at least 14 heterotypic validly described synonyms at the specific level and several more as variants or sub- varieties for the annual *Brachypodium distachyon sensu lato*. Some of the synonymized names were described as annual species independent from *B. distachyon* (e.g., *Festuca monostachyos* Lamarck (1788: 461), *Festuca rigida* Roth (1797: 12), *Triticum brevisetum* Candolle (1813: 153), *Bromus pentastachyos* Tineo (1817: 4), *Brachypodium megastachyum* Besser ex Schultes & Schultes f. (1827: 3), *Brachypodium macrostachyum* Besser ex Schultes & Schultes f. (1827: 3), among other), although this author subsumed them within *B. distachyon sensu stricto* as he considered they were part of the large phenotypic variation of a single species *B. distachyon*.

Catalán *et al*. (2012) could not analyze molecularly the type LINN 93.48 nor the heterotypic synonyms, due to prohibition for destructive sampling of types in their hosting herbaria, but analyzed statistically some morphological features of LINN 93.48 together with those of representative samples from the three species. However, due to the ambiguous classification of LINN 93.48 in the discriminant analysis, falling in an intermediate position between the *B. distachyon* cluster and *B. stacei* in the bidimensional DA plot, these authors chose a Bd21 (2n=10) accession specimen as an epitype for *B. distachyon*, according to Art. 9.9 of the Shenzhen Code or *ICN* (Turland *et al*. 2018 [Art. 9.7 in the Vienna Code, see McNeill *et al*. 2005]). They also selected holotype specimens for the newly described *B. stacei* and *B. hybridum* species from the respective ABR114 (2n=20) and ABR113 (2n=30) accessions’ materials (Catalán *et al*. 2012).

Although subsequent studies enlarged the morphological characterization of *B. distachyon*, *B. stacei*, and *B. hybridum* with more diagnostic quantitative and qualitative morphoanatomical traits (Catalán *et al*. 2016b, López-Álvarez *et al*. 2017), and the generation of reference genomes for *B. stacei* and *B. hybridum* (Gordon *et al*. 2020, Scarlett *et al*. 2023, Mu *et al*. 2023a, 2023b), the nomenclature of the three annual species of the *Brachypodium distachyon* complex is not yet established. Namely, a revision of the heterotypic synonyms of the annual *Brachypodium* species is still lacking. In this paper we have analyzed statistically morphological diagnostic traits that could be measured in the type specimens of 11 validly published names (*Brachypodium distachyon* (≡ *Bromus distachyos*), *B. hybridum*, *B. macrostachyum, B. megastachyum*, *B. stacei*, *B. pentastachyos*, *Festuca monostachyos*, *F. rigida*, *Trachynia platystachya* (Balansa ex Coss.) Scholz in Greuter & Raus (1998: 173) [≡ *Brachypodium distachyon* var. *platystachyon* Balansa ex Coss. in Cosson & Durieu (1855: 192)], *Triticum schimperi*, and *T. subtile* Fischer *et al*. (1845: 59) and have conducted a thoroughly nomenclatural revision. Unfortunately, assessing the chromosome number or ploidy level of annual *Brachypodium* herbarium specimens is a challenge, as these diagnostic traits are correlated to microscopical anatomical features (e. g., leaf stomatal guard cell length, pollen grain length) (Catalán *et al*. 2012), that are extremely difficult (or almost impossible) to measure through non-destructive sampling of type specimens. Similarly, fully diagnostic molecular data can’t be retrieved from them as destructive study of valuable herbarium types is not a recommended practice and is even forbidden due to some herbarium conservation policies (Rabeler *et al*. 2019, Rosselló *et al*. 2021, Davis *et al*. 2024). According to our results, a proposal for the conservation of the names *B. hybridum* and *B. stacei* is needed and will be presented. Our study is a further step in our contribution to completing the nomenclature of the grass model genus *Brachypodium* (Catalán *et al*. 2012, Fabado & Ferrer-Gallego 2021, Ferrer-Gallego & Martínez- Labarga 2022a, 2022b, Ferrer-Gallego & Fabado 2023).

## Materials and methods

### Search and selection of names

For the selection of the traditionally synonymous names of *Brachypodium distachyon sensu lato*, we relied primarily on the work of Schipmann (1991) and on the current nomenclatural databases POWO and WFO. In total, we reviewed the protologues of 16 names, in addition to the accepted names *B. distachyon*, *B. hybridum*, and *B. stacei*. Of these 16 names, original type material was found for 9, allowing us to perform the taxonomic and nomenclatural analyses. For the remaining names, we attempted to interpret the descriptions and searched for any illustrations that could help link them to one of the annual *Brachypodium* species or to other species.

### Sampling and morphological and statistical analyses

The sampling was designed to comparatively analyze the taxonomic assignments of the herbarium types of the basionym and validly published heterotypic synonyms of *Brachypodium distachyon sensu lato* with the herbarium types of the currently recognized species *B. distachyon*, *B. stacei*, and *B. hybridum*, and a representative sampling of 50 other specimens of the three species (Table 1). The lectotype of *B. distachyon*, specimen LINN 93.48, has been reanalyzed in the present study to better re-evaluate its possible taxonomic assignment. In total, 63 samples were incorporated in the morphological discriminant analysis (DA): the type specimens of the currently recognized *Brachypodium distachyon* (Iraq: Salah ad Din, 4 km from Salahuddin; epitype: MA833764, isoepitype: JACA-R298981), *B. stacei* (Spain: Balearic isles: Formentera; holotype: MA 833765; isotype: JACA-R298982) and *B. hybridum* (Portugal: Lisboa; holotype: MA 833766, isotype: JACA-R298983), eight additional herbarium types of heterotypic synonyms (*Brachypodium macrostachyum*, *B. megastachyon*, *Bromus pentastachyum*, *Festuca monostachyos*, *F. rigida*, *Trachynia platystachya*, *Triticum schimperi*, *T. subtile*), and another fifty specimens of representative samples of these taxa (*B. distachyon*: 16; *B. stacei*: 16; *B. hybridum*: 18), with the aforementioned LINN 93.48 (Table 1). The taxonomic identity of all non-type samples was previously confirmed by chromosome counting, molecular barcoding, and/or genomic data (Catalan *et al*. 2012, Catalan *et al*. 2016b, Lopez-Alvarez *et al*. 2017, Gordon *et al*. 2020, Campos *et al*. 2024).

**TABLE 1.**
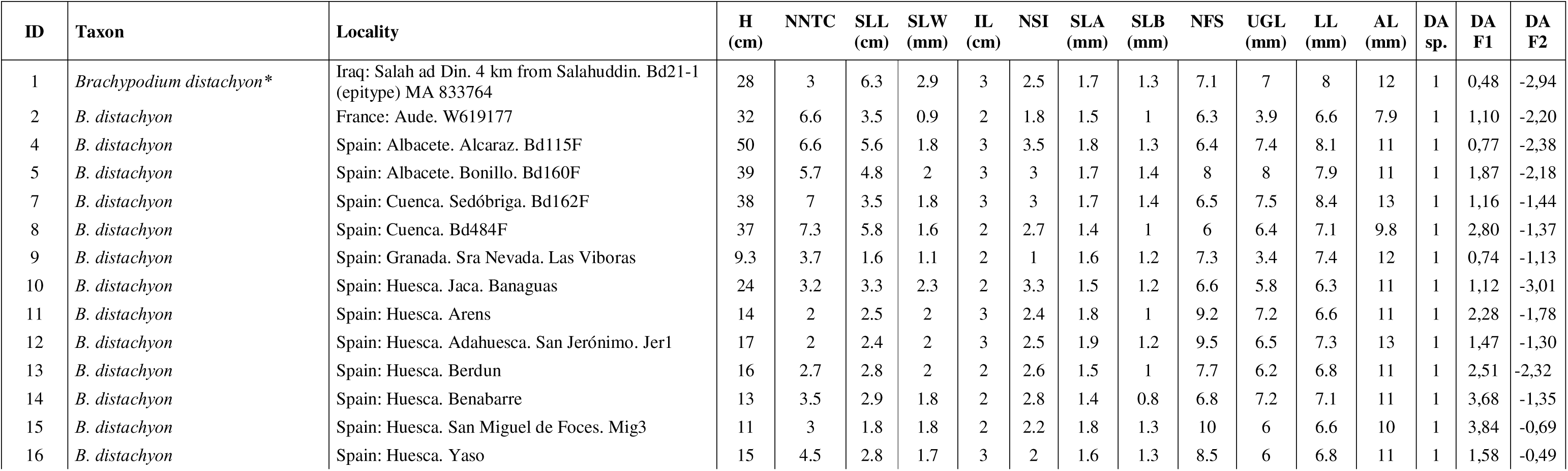
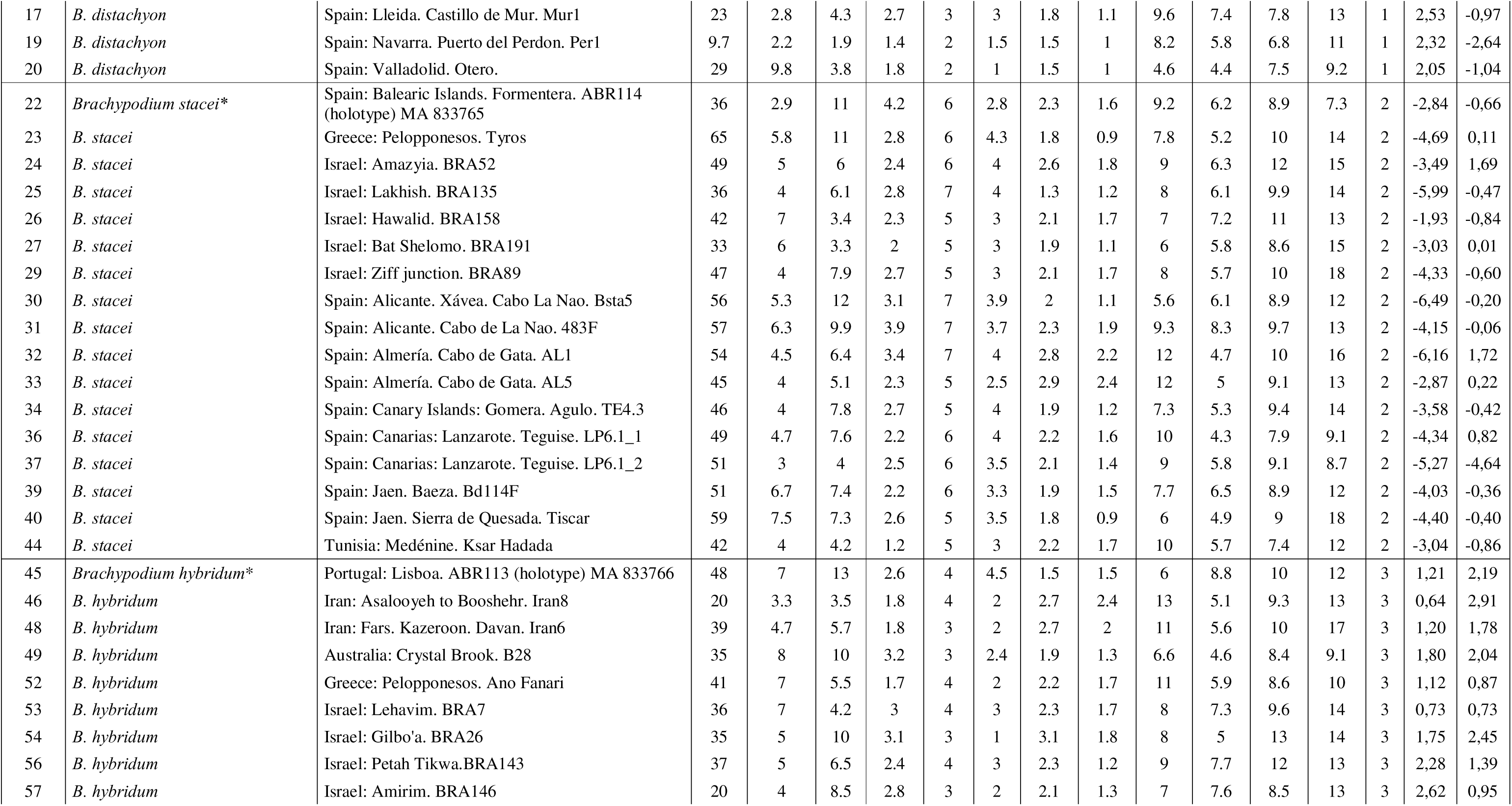
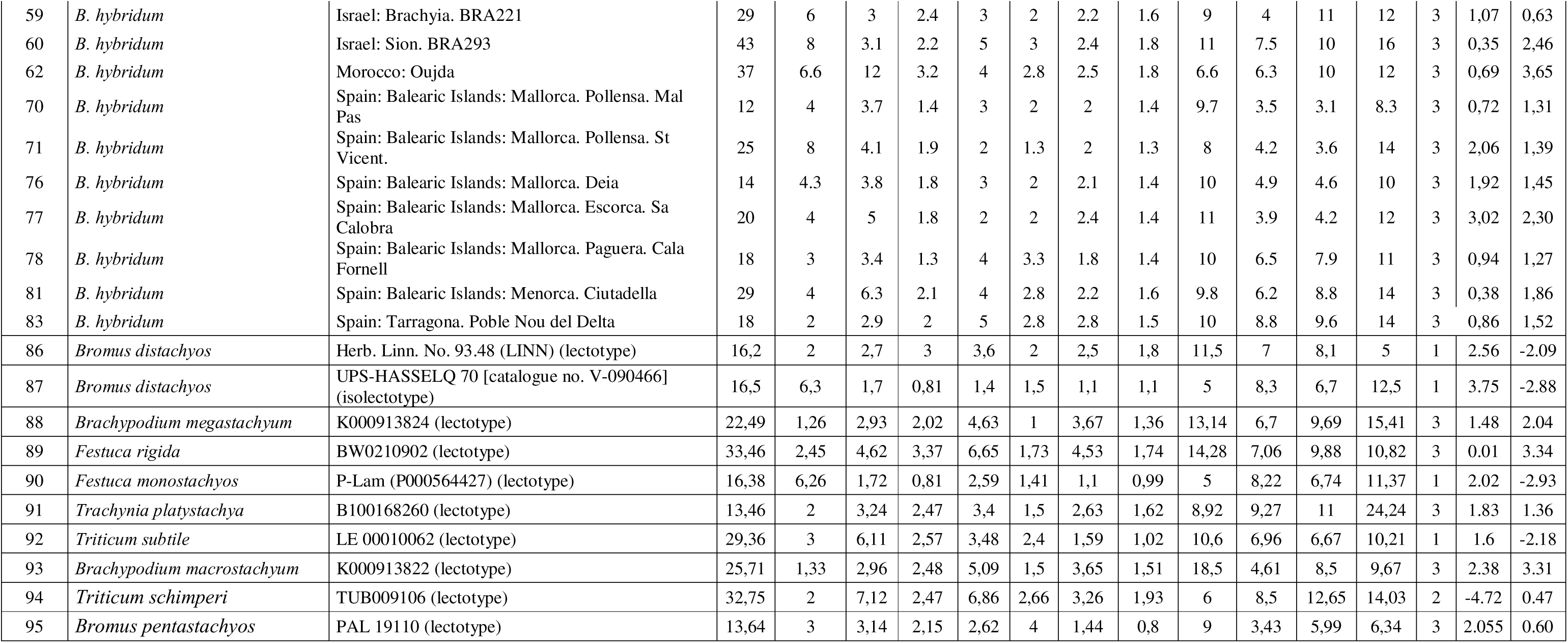
Sources of the three *Brachypodium distachyon*. *B. stacei* and *B. hybridum* type samples, of 50 wild samples confirmed taxonomically and genomically to belong to each annual *Brachypodium* species. and of nine heterotypic synonyms’ type samples that were included in the morphological discriminant analysis. Asterisks indicate *B. distachyon*. *B. stacei* and *B. hybridum* type samples and diamonds type samples of the heterotypic synonyms under study. Measurements of morphological variables correspond to averaged values from up to 5 measurements per trait and individual (and from up to 10 individuals per population for the wild samples). ID: numerical identity of each sample. Abbreviation of variables: H (Plant height). NNTC (Number of nodes of tallest culm). SLL (Second leaf length from the base of the plant). SLW (Second leaf width). IL (Inflorescence length). NSI (Number of spikelets per inflorescence). SLA (Spikelet total length. without awns). SLB (Spikelet length from the base to the apex of the fourth lemma. without awns). NFS (Number of flowers per inflorescence). UGL (Upper glume length). LL (Lemma length from the basal floret). AL (Awn length. the longest within the spikelet). The DA sp. column indicates the classification of samples to *B. distachyon* (1). *B. stacei* (2) or *B. hybridum* (3) species according to the results of the discriminant analysis. Columns DA F1 and DA F2 show the values of Function 1 and Function 2 obtained for each sample in the discriminant analysis (their projections are shown in the two-dimensional DA graph in Fig. 1).

We analyzed a total of 12 quantitative and discrete morphological characters that could be measured in the type specimens studied and that were employed to separate and identify the three species of the *B. distachyon* s. l. complex in our previous work (Catalán *et al*. 2012, López-Álvarez *et al*. 2017). These included (Plant) Height (H) (cm), Number of Nodes of Tallest Culm (NNTC), Second Leaf Length (SLL) (cm), Second Leaf Width (SLW) (mm), Inflorescence Length (IL) (cm), Number of Spikelets per Inflorescence (NSI), Spikelet Length (total, without awns; SLA) (mm), Spikelet Length (from base to 4^th^ lemma, without awns; SLB) (mm), Number of Flowers per Inflorescence (NFI), Upper Glume Length (UGL) (mm), Lemma Length (LL) (mm), and Awn Length (AL) (mm). Morphological characters were measured with a hard ruler under a dissecting microscope on herbarium samples and with Annotate-on v.1.9.56 tool (RECOLNAT-ANR-11- INBS-0004) for magnified type specimens imagens.

Following the procedures of statistical analysis performed with quantitative and discrete morphological characters for the classification of newly studied samples in our previous studies of the annual *Brachypodium* species (Catalán *et al*. 2012, Lopez-Alvarez *et al*. 2017), we performed a Discriminant Analysis (DA, cross-validation) with all variables (12) and samples (62) under study to determine the group of belonging with greater probability of the samples (Legendre & Legendre 1998) and the assignment of the new type specimens to their respective optimal group. We used as reference samples for the group of each species the respective type specimens of the current species (*B. distachyon*: Bd21; *B. stacei*: ABR114; *B. hybridum*: ABR113) (Table 1). The identification of the more discriminating variables was done through Fisher’s coefficient at the significant threshold value of 0.05; the posterior probability of classification of each sample and of each discriminant function were calculated through Wilks’ Lambda values (a Wilks’ Lambda value closer to zero indicated a better discrimination between the predefined groups). The DA was run in SPSS v. 29.0. We pre-assigned the samples of the newly studied type specimens to the currently recognized annual *Brachypodium* species according to their morphological proximity, based on the 12 quantitative and discrete traits studied, and searched for best assignments in the DA output, where the classification of all samples was 100% correct.

### Typification of the names

The typifications discussed and included in this work are based on the analysis of the protologues of the names, the examination of relevant literature and on the study of the specimens conserved in several herbaria. Herbarium acronyms are cited according to Thiers (2025 [continuously updated]), some of which are available as virtual herbaria on-line. In typifying names of taxa, we strictly followed the International Code of Nomenclature for algae, fungi, and plants (Turland *et al*. 2018). The identity of the designated types has been verified with the current use of their respective names. The typified names are followed by their homotypic synonyms (indicated with the symbol ≡) and the heterotypic synonyms (indicated with the symbol =). The names in current use are set in bold italics typeface.

## Results and Discussion

### Taxonomic Analysis

Taxonomic analysis of the type specimens of eight names corresponding to heterotypic synonyms of *Brachypodium distachyon* s. l. and the statistical discriminant analysis based on 12 phenotypic traits have provided congruent results. The DA classification performed on the 62 samples of individuals of *B. distachyon*, *B. stacei* and *B. hybridum* plus eleven type gatherings (eight of heterotypic synonyms, two of newly described species and the epitype of *B. distachyon*) using the data of the 12 morphological traits analysed resulted in the correct classification of all samples into their respective predefined groups (Table 1). All the *B. distachyon*, *B. stacei* and *B. hybridum* non- type samples were correctly classified (100%) into their respective species; furthermore, the DA assigned the type specimens of *B. distachyon* (lectotype and epitype), *Festuca monostachyos* and *Triticum subtile* to group 1 (*B. distachyon*), the type specimens of *B. stacei* (holotype) and *Triticum schimperi* to group 2 (*B. stacei*), and the type specimens of *B. hybridum* (holotype), *Brachypodium distachyon* var. *platystachyon* (≡ *Trachynia plastystachya*), *B. macrostachyon*, *B. megastachyon*, *Bromus pentastachyos* and *Festuca rigida* to group 3 (*B. hybridum*) (Figure 1, Table 1).

**FIGURE 1.**
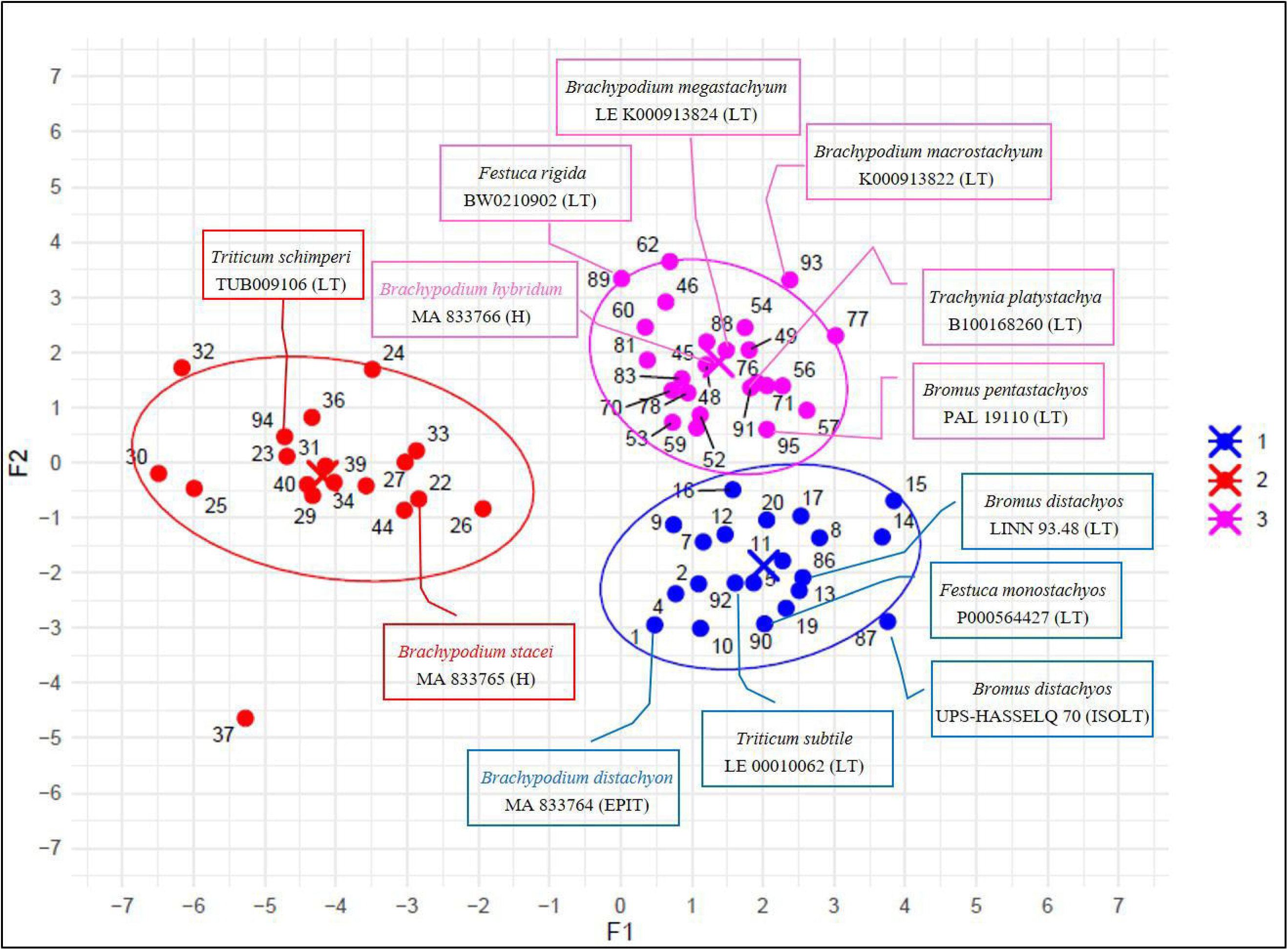
Two-dimensional DA scatterplot of samples of *Brachypodium distachyon* (group 1, blue), *B. stacei* (group 2, red), and *B. hybridum* (group 3, purple) and of type specimens assigned to each group based on averaged values of samples analyzed for 12 quantitative and discrete morphological traits in 63 samples (see text and Table 1). The first and second canonical discriminant functions explained 71.5% and 28.5% of the interspecific taxonomic variation, respectively. Crosses indicate the group centroid. Ellipses represent the 95% probability of group membership. Numerical identities of samples correspond to those in Table 1. The types studied are indicated in the plot.

In the two-dimensional DA plot constructed with the two functions, *B. stacei* clustered separately from *B. distachyon* and *B. hybridum* along the first axis of the plot, which explained 71.5% of the total variance, whereas the last two species separated along the second axis, which explained 28.5% of the variance (Figure 1). The DA confirmed that the morphological separation of the three species is supported by four characters (IL, H, SLA, SLB) (Table S1). Wilks’ Lambda values of the first and second discriminant functions were, respectively, 0.033 and 0.279. The lowest Wilks’ Lambda value obtained for the first discriminant function that separated the *B. stacei* from the other species indicated a greater morphological differentiation of this diploid from its congeners, while the low value obtained from the second discriminant function also supported the morphological distinction of *B. distachyon* from *B. hybridum*. Interestingly, our DA results differ slightly from those of previous studies in the greatest morphological separation of *B. stacei* from the rest (Figure 1) while those of Catalan *et al*. (2012) and Lopez-Alvarez *et al*. (2017) showed the greatest morphological differentiation of the allotetraploid *B. hybridum* from its diploid progenitor species. This may be a consequence of the lower number of quantitative and discrete characters used in our current DA study (12 phenotypic traits) than in those of Catalan *et al*. (2012) and Lopez-Alvarez *et al*. (2017) (15 phenotypic traits) due to the impossibility to measure Caryopsis Length (CL) and, crucially, the highly discriminant microanatomical (Stomata) Leaf Guard Cell Length (LGCL) and Pollen Grain Length (PGL) traits in our non-destructive study of the type specimens. These results also indicate that for our well represented sampling of the three species, *B. stacei* is the most distinct species of the trio regarding the twelve analysed morphological characters (H, NNTC, SLL, SLW, IL, NSI, SLA, SLB, NFI, UGL, LL, AL) (Table 1), and especially with respect to the four traits that support the main functions of our DA (IL, H, SLA, SLB) (Table S1).

We consider the current DA to be significantly more robust than previous analyses (Catalan *et al*. 2012), primarily due to the better representation of molecularly corroborated specimens of the three species (*B. distachyon*, *B. hybridum*, and *B. stacei*), including the reassessment of Iranian samples previously misclassified as pertaining to *B. stacei* and recently identified as belonging to *B. hybridum* (Campos *et al*. 2024), that cover a large portion of their natural distributions, as well as areas of recent introduction (Table 1). However, we acknowledge the difficulties—and potential errors—involved in measuring 12 quantitative characters (many of which are of small dimensions) and qualitative traits from images of old herbarium specimens. Despite these challenges, the reliability of these measurements has been demonstrated in other studies (Borges *et al*. 2020). This issue may be reflected in the different classification of the *Bromus distachyos* lectotype LINN 93.48, which was considered intermediate between *B. distachyon* and *B. stacei* in Catalán *et al*. (2012), based on only one sample of *B. stacei*. In our current study, however, with 17 or more samples per species (Tables 1, S1, Figure 1), it was assigned to *B. distachyon*.

We have adopted a consensus strategy for assigning type specimens to extant species and have successfully classified all types, with the assignment supported by the DA data.

### Nomenclatural treatment and typification of the names

We examined the nomenclature of each of the proposed names once considered heterotypic synonyms of *B. distachyon sensu lato*, grouping them into the three recognized annual species of *Brachypodium* (*B. distachyon*, *B. hybridum*, and *B. stacei*) based on our taxonomic revision and the DA output (Figure 1, Tables 1, S1, and 2).

***Brachypodium distachyon*** (L.) Palisot de Beauvois (1812: 155, 156, 101)

The traditional concept and current use of the name *B*. *distachyon* has been applied to a stiffly erect annual plant up to (2–)6–15(–56) cm high, leaf blades of culms (1–)3–7(–8.5) cm × (0.25–)1.5–2(– 4) mm, erect, flat, unbendable, straigth, bright green, usually villose abaxially and adaxially, sheaths glabrous to densely villose abaxially, panicle with 1–2(–4) spikelets, spikelets (9.2–)14–15(–24) mm, and lemmas (5.2–)7–8(–11) mm, 7-veined, awned, with awn (5–)10–11(–15.2) mm (see Catalán *et al*. 2012, 2016a, 2016b). The lectotype of *Bromus distachyon* was designated by Schippmann & Jarvis (1988) as “neotype” from a specimen preserved at LINN 93.48 (image available at https://linnean-online.org/982/). However, Jarvis (2007) mentioned that the specimen is a Fredric Hasselquist collection, closely associated with the thesis in which the binomial appeared (Stearn 1957), and therefore this sheet is part of the original material and should be treated as a lectotype under Art. 9.8 *ICN* (Turland *et al*. 2018). There is also an isolectotype preserved at UPS- HASSELQ 70 [catalogue no. V-090466] (see Jarvis 2007). The sheet at UPS bears a plant, with leaves and inflorescences, and is annotated “agypt” by Fredric Hasselquist at the base of the specimen.

According to Catalán *et al*. (2012, 2016a) the type could not be confidently and unambiguously assigned to any of the three species of the *B. distachyon* complex due to their intermediate morphological diagnostic characters. Specifically, the specimen LINN 93.48 shows intermediate phenotypic features between the 2*n* = 10 (*B*. *distachyon*) and 2*n* = 20 (*B*. *stacei*) taxa. In addition, it was impossible to study this specimen cytogenetically or molecularly, thus precluding the analysis of its chromosome number or molecular barcodes, characters also used to identify and discriminate the species.

As the identification of the lectotype cannot be critically and precisely attributed to *B*. *distachyon* in the current usage and concept of this name, Catalán *et al*. (2012) tried to remedy this situation by designating an epitype for *B. distachyon* from a molecularly tested specimen from Iraq that corresponds to the assembled and annotated Bd21 (2*n* = 2*x* = 10) reference genome (see Catalán *et al*. 2012, 2016a).

The specimen selected as epitype, preserved at MA (with barcode MA 833764), is a complete and well-preserved specimen that corresponds with the current usage of the name and matches the traditional concept of *B*. *distachyon* (see, e.g., Schippmann 1991, Catalán *et al*. 2012, 2016a). This material was obtained from the same near-isogenic-line (NIL) of the material used to build the reference genome of *B. distachyon* (accession Bd21; see IBI 2010). The specimen shows the diagnostic features described above, with available duplicate specimens at JACA (code JACA- R298981), JE (barcode JE00013261), and K (barcode K000743625), and molecular and cytogenetic data. This specimen will maintain the name of the most commonly investigated species of *Brachypodium*, which is today an important model plant for genomic studies of cereal and grass biofuel crop relatives.

In the present work, the taxonomic assignment of the LINN 93.48 sheet has been re-evaluated, together with the UPS-HASSELQ 70 [catalogue no. V-090466] isolectotype. If in the previous analysis (Catalán *et al*. 2012) it was only possible to compare the samples with a single specimen of *B. stacei*, in the current analysis the representation of the three species has been much higher, which has allowed us to place the LINN 93.48 and UPS-HASSELQ 70 [catalogue no. V-090466] sheets within the range of variability of *B. distachyon* sensu stricto.

*Festuca monostachyos* Lamarck (1788: 461)

Lamarck’s protologue of *Festuca monostachyos* (1788: 461), cited as “13. Fétuque à un épillet” in the *Encyclopédie Méthodique*, consists of a morphological description in Latin: “Festuca spicula unica terminali, aristis longis, foliis margine ciliatis. N. [Nobis]”, followed by a complete description of this species in French. Furthermore, Lamarck (1788) added “Cette plante croît sur la côte de Barbarie, & nous a été communiquée par M. l’Abbé Poiret (v.s.)” [This plant grows on the coast of Barbary, and was communicated to us by M. l’Abbé Poiret)].

We have located a specimen at P, with barcode P00564427, which is part of the original material used by Lamarck to describe his species. The sheet bears three plants, with leaves and spikelets, and an original label annotated by Lamarck “Festuca monostachyos. lam. / dict. / Crôit sur la côte de barbarie, selon / M. l’abbé Pourret”. This specimen is designated is this paper as the lectotype of the name *Festuca monostachyos*. According to our DA morphological classification, the position of specimen P00564427 allows us to consider the name *Festuca monostachyos* as conspecific with *B. distachyon* and therefore a heterotypic synonym of the Linnaean name.

***Triticum subtile*** *Triticum subtile* Fischer, Meyer & Avé-Lallemant (1845: 59)

*Triticum subtile* was published by F.E.L. von Fischer, C.A. Meyer and J.L.E. Avé-Lallemant (in Index Seminum [St. Petersburg (Petropolitanus)] 10: 59. 1845) with a complete description in Latin. The protologue, numbered “2669” also includes the phrase “Trachynia subtilis Hort.

Genuens., in Ind. sem. hort. Univ. Lipsiens. p. ann. 1842”. There are two relevant specimens at LE, with barcodes LE00010062 and LE00010063. This material was cultivated in the *Hortus Botanicus Imperialis Petropolitanus* and collected by Carl Anton Meyer in 1844. The sheet barcoded LE00010062 bears a good specimen and two handwritten labels: 1) “Typus! / Triticum subtile Fisch. et Mey. / Ind. Sem. Hort. Petrop. X (1845), 59 / “Trachynia subtile Hort. Genuen. in Ind. sem. hort. / Univ. Lipsiens. p. ann. 1842”, and 2) “Trachyna subtilis / H. Genuens. / Sem. Kunze / Cult. in h. b. Petropolit. / 1844 / Meyer”. The sheet barcoded LE00010063 bears also a good specimen and two labels, annotated as: 1) “subtile / C.A. Meyer”; and 2) “Trachynia / subtilis / m. Kunze h. / Genean.”. Both specimens are well-preserved and complete. The specimen LE00010062 is designated in this work as the lectotype of the name.

According to our results, the names *Bromus distachyos* and *Triticum subtile* could be considered conspecific. Therefore, the latter is a new heterotypic synonym of *B*. *distachyon*.

***Brachypodium distachyon*** (L.) Palisot de Beauvois (1812: 101, 155, 156. 1812 [‘*distichyos*’, ‘*distachyum*’])

≡ *Bromus distachyos* Linnaeus (1756 (10 March): 13; 1756 (2 June): 8)

**Lectotype** (designated by Schippmann & Jarvis (1988) as “neotype”): Herb. Linn. No. 93.48 (LINN). **Isolectotype**: UPS-HASSELQ 70 [catalogue no. V-090466] (Figure 2). **Epitype** (designated by Catalán *et al*. 2012): Iraq, Salah ad Din, 4 km from Salahuddin, in the road to Mosul. Col. No. K1202. USDA PI 254867, Bd21 (inbred line, obtained from seeds cultivated at Aberystwyth University, 30 October 2010, *Luis A.J. Mur s.n.*, used to generate the reference genome of *Brachypodium distachyon*) (MA barcode MA 833764; **isoepitypes**: JACA code JACA-R298981, JE barcode JE00013261, K barcode K000743625).

**FIGURE 2.**
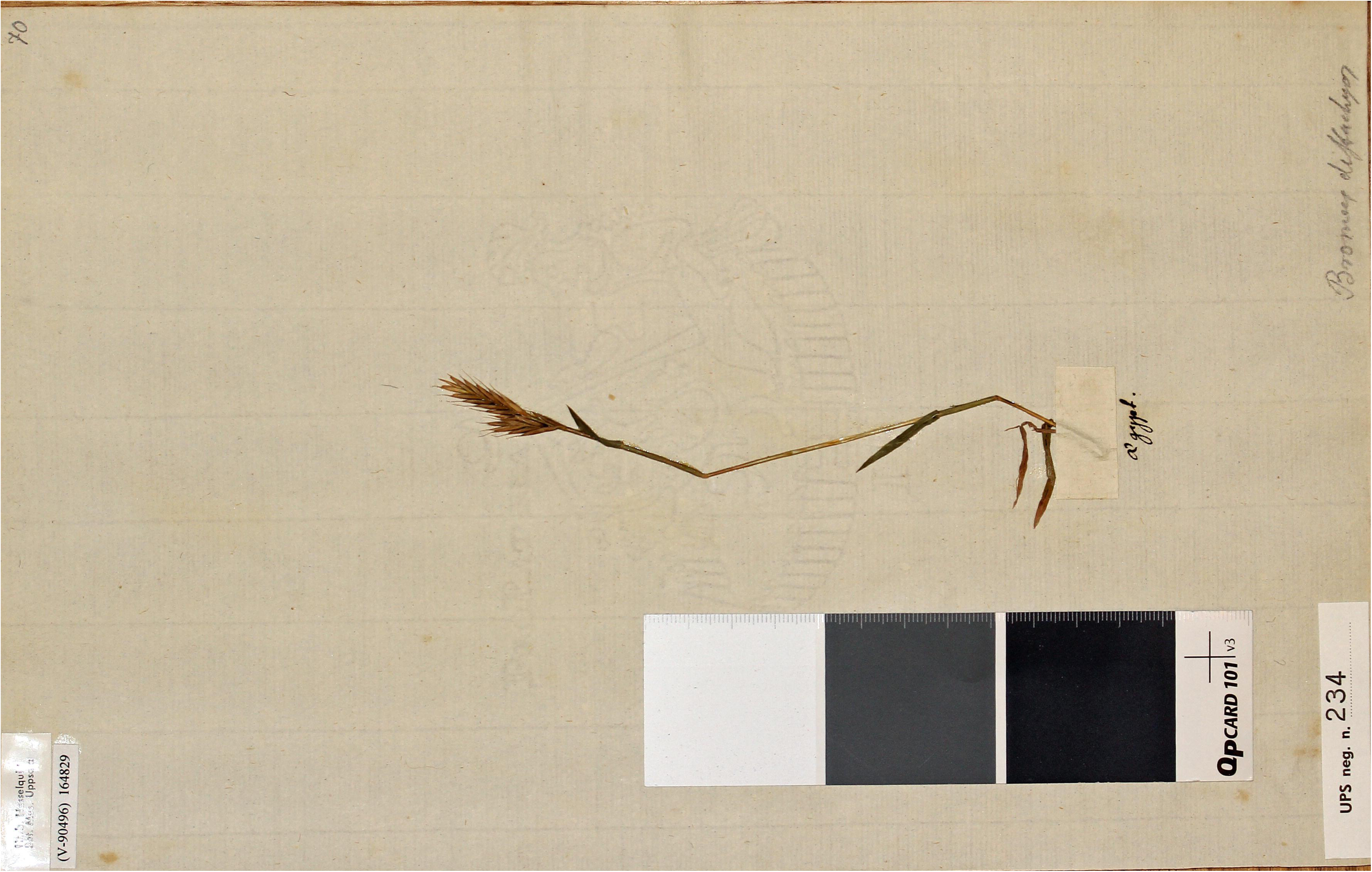
Isolectotype of *Brachypodium distachyon*, UPS-HASSELQ 70 [catalogue no. V- 090466]. Image by courtesy of the herbarium UPS, reproduced with permission.

= *Festuca monostachyos* Lamarck (1788: 461), ***syn. nov*.**

**Lectotype** (**designated here**): Crôit sur la côte de Barbarie, selon M. l’abbé Pourret, s.a., s.d., (P barcode P00564427) (Figure 3).

**FIGURE 3.**
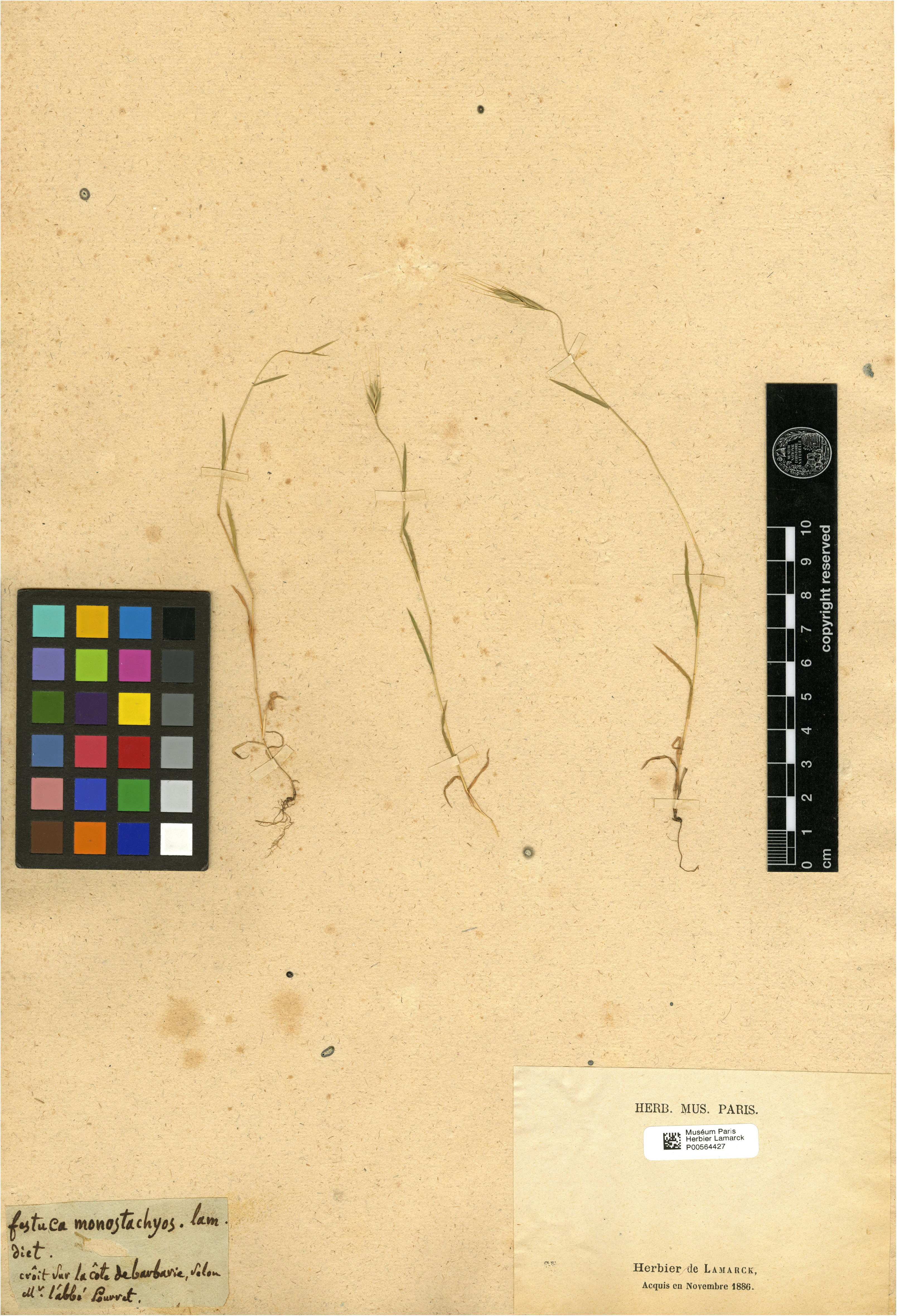
Lectotype of *Festuca monostachyos* Lam., P barcode P00564427. Image by courtesy of the herbarium P, reproduced with permission.

= *Triticum subtile* Fischer, Meyer & Avé-Lallemant (1845: 59), ***syn. nov*.**

**Lectotype** (**designated here**): Russia, 1844, *C.A. Meyer s.n*., LE barcode LE00010062 (Figure 4).

**FIGURE 4.**
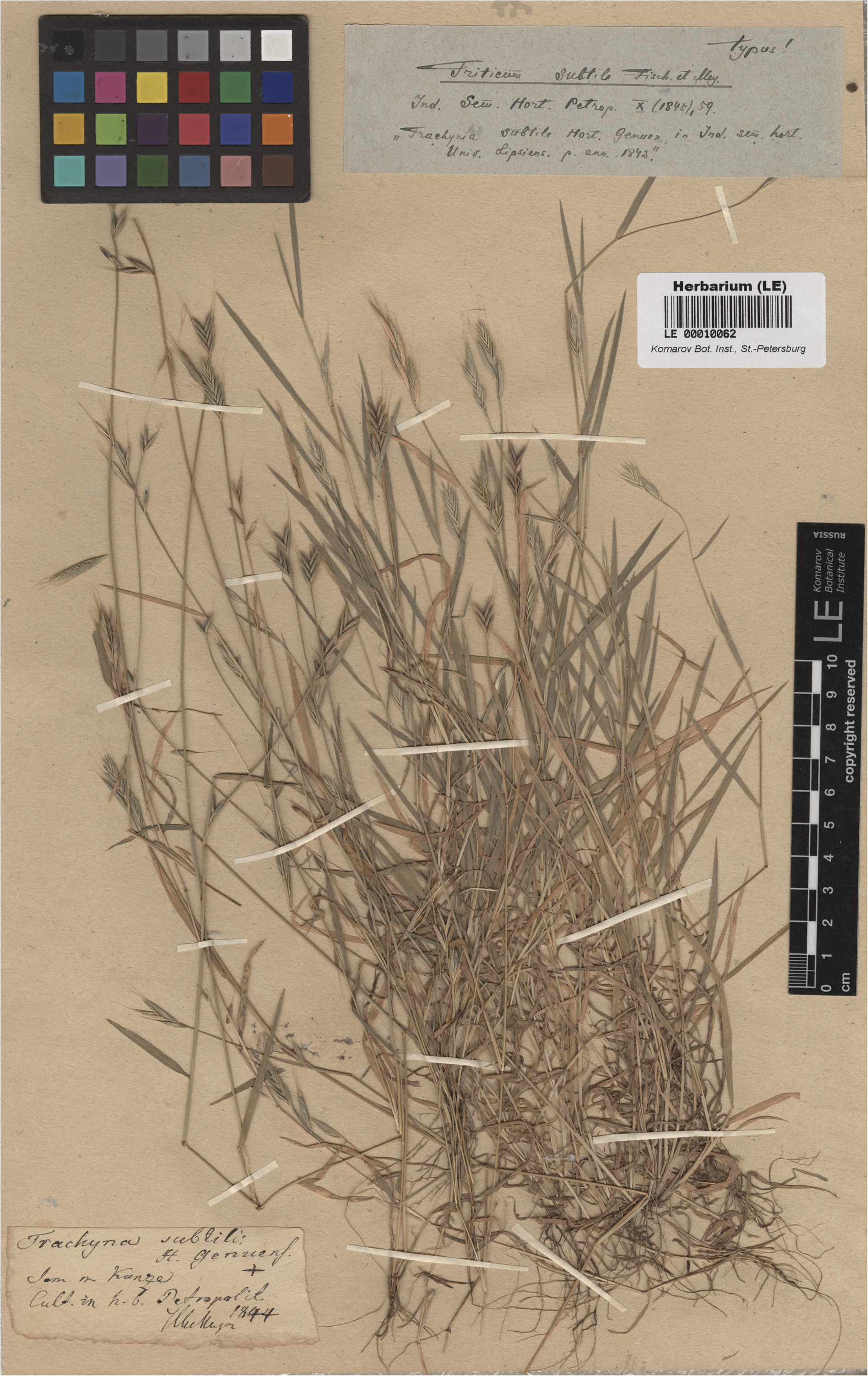
Lectotype of *Triticum subtile* Fisch., LE barcode LE00010062. Image by courtesy of the herbarium LE, reproduced with permission.

Isolectotype: LE00010063.

***Brachypodium hybridum*** Catalán, Joch.Müll., Hasterok & G. Jenkins (2012: 402 [18 of 21])

*Brachypodium hybridum* is a species native to the circum-Mediterranean region (also recorded in SW Asia: Afghanistan, Armenia, Iran, Iraq, Kuwait), and Macaronesia (Spain: Canary Islands: Fuerteventura, Gomera, Lanzarote, Tenerife) (Catalán *et al*. 2012, López-Álvarez *et al*. 2015, 2017, Mu *et al*. 2023b). The species has been introduced in other regions of central Europe, western- northern America (California), southern America (Uruguay and Argentina), South Africa, and Oceania (Australia and New Zealand) (Jenkins *et al*. 2003, Garvin *et al*. 2008, Bakker *et al*. 2009, Catalán *et al*. 2012, 2016b) and is considered an invasive plant in California and Australia (López- Álvarez *et al*. 2017). *B. hybridum* can grow in sympatry with (or more likely near) its *B*. *stacei* or *B*. *distachyon* progenitor species in some Mediterranean spots (Shiposha *et al*. 2020).

The current concept and use of the name *B*. *hybridum* is applied to an erect or spreading annual plant up to (3.5)30–40(78) cm high, leaf blades of culms 7–8(16) cm × (0.7)2–3(4.3) mm, glabrous to pubescent abaxially, with panicle with 3(–6) spikelets, spikelets (10)18–24(41) mm, and lemmas (3)8–10(12.9) mm, with awn (6)11–12(18.9) mm (see Catalán *et al*. 2012, 2016a, 2016b).

***Festuca rigida*** Roth (1797: 12)

The protologue of *Festuca rigida* Roth (1797: 12) consists of the name *Festuca rigida* numbered “3”, followed by a diagnosis “spica terminali oblonga, subcompressa, seminibus ovato-oblongis aristatis imbricatis, culmo incrassato”, a complete and extensive description in Latin, and the provenance “Habitat in *Hispania*”. The protologue also contains two comments regarding the variability of the taxon and the differences with *Brachypodium distachyon* (mentioned as *Festuca distachyo*).

Albrecht Wilhelm Roth (1757–1834)’s herbarium is preserved today at B and B-Willdenow, among other herbaria (e.g., W) (see Stafleu & Cowan 1983). In the B-Willdenow herbarium there are three relevant sheets, with barcodes B-W 02109-01 0, B-W 02109-02 0, and B-W 02109-03 0, within the Willdenow folder: B-W 02109 (image available at https://ww2.bgbm.org/Herbarium/specimen.cfm?Barcode=BW02109000). The sheet barcoded B-W 02109-01 0 bears four plants, with leaves and inflorescences, and is annotated as “*F*. *rigida*. / 1” on the upper right corner and “W.” [Willdenow] at the base of the sheet. The sheet B-W 02109-02 0 bears two complete and well developed plants, with leaves and inflorescences, and is annotated as “*F*. *rigida*. / 2” on the upper right corner and “W.” at the base of the sheet. Finally, the sheet B-W 02109-03 0 bears only a plant, which shows a culm with two leaves and four spikelets, and is annotated “*F*. *rigida*. / 3” on the upper right corner and “W.” at the base of the sheet, the sheet contains also a handwritten label “*Festuca* / *rigida*”. In the herbarium of the Naturhistorisches Museum Wien (W), we have also found a relevant specimen, with the barcode W-0053543, consisting of a plant with leaves and inflorescence, accompanied by three handwritten labels in the lower right corner. The lowest one reads: “Herb. Wulfen. / Triticum / hispanicum / Rothio Festuca rigida”, probably written by F.X. Wulfen (see Burdet 1979). Above this there is another label that reads: “Festuca rigida Roth. / Species nova Hispanica. / quidni potius Triticum?”. On these two labels we find a review label by MW van Slageren (RBG Kew; 30/VII/2014) indicating that the specimen belongs to the *Brachypodium distachyum* group. Although it could be a sheet studied by Roth himself, we are not absolutely certain that this is the case, and therefore have discarded this specimen as original material. We have not been able to locate any further original material for the name *F*. *rigida*.

Among these specimens, we designate the specimen with barcode B-W 02109-02 0, as the lectotype of the name *Festuca rigida*. This specimen is the most complete and informative original material, it matches the protologue and the traditional concept, and corresponds to the species as currently delimited.

According our data, the analysed specimen at B (B-W 02109-02 0), designated as the lectotype of *Festuca rigida*, can be identified as belonging to *Brachypodium hybridum*, and these names should be therefore treated as synonyms.

*Bromus pentastachyos* Tineo (1817: 4)

*Bromus pentastachyos* was described by Tineo (in Rar. Sicil. 1: 4. 1817) and the protologue consist in a complete description in Latin, followed by word “Annuus” (i.e., annual plant) and “*Hab. l.c.*”, the same place as the preceding species, “prope Agrigentum” (Sicily, Italy). The lectotype was indicated by Steinberg (1981) and supported by Schippmann (1991) based on herbarium specimen from Palermo, PAL19110.

This is a scarce but complete sheet, consisting of a plant with leaves and inflorescences, accompanied by two labels, one next to the plant with the inscription “HOLOTYPUS / Bromus pentastachyos Tin.”, and another handwritten one, on the lower left hand side, where it is written “Brachyp. distachion. var. 4-5. / stachyum / var. minor / B. pentastachyos nob.”

This is a name used by some floras to designate plants independent of *B. distachyon* through the combination *Trachynia pentastachya* (Tineo) Tzvelev (1975: 81, Takhtazhan 2006), although, according to our data, the analysis of the specimen PAL19110 can be identified as belonging to *Brachypodium hybridum*.

***Brachypodium macrostachyum*** Besser ex Schultes & Schultes f. (1827: 3)

The name *Brachypodium macrostachyum* Besser (in Mant. 3 (Schultes & Schultes f.) 3: 651. 1827) was published with a brief description followed by the annotation “*Besser in litt*.”, and the number “n. 11b”. Schippmann (1991: 178) indicated that the “typus” is preserved at K. We have located a specimen at K (barcode K000913822), which is part of the original material. The sheet at K bears four well-preserved and complete plants and a revision label annotated by Schippmann indicating that it is the type.

Based on our results, the names *Brachypodium hybridum* and *Brachypodium macrostachyum* could be considered conspecific. Therefore, *B. macrostachyum* is a new heterotypic synonyms of *B*. *hybridum*.

***Brachypodium megastachyum*** Besser ex Schultes & Schultes f. (1827: 3)

Another name which is relevant for the nomenclature of *B. hybridum* is *B. megastachyum* Besser (in Mant. 3 (Schultes & Schultes f.) 3: 651. 1827): 196. The name *Brachypodium megastachyum* was published with a brief description, the number “n. 11a”, and the comment “Forsan varietas multiflora, culturâ orta Br. *distachyi*, cujusque sub nomine quoque teneo. *Besser*.”.

We have located three relevant specimens at K (barcode K000913824) and LE (barcodes LE00009972 and LE00009971), which are part of the original material. The specimen at K was designated by Schippmann (1991: 178) as the type of the name. According to our data, this specimen is included in the group of *Brachypodium hybridum*.

***Brachypodium distachyon*** var. ***platystachyon*** Balansa ex Coss. in Cosson & Durieu (1855: 192)

Cosson (in Cosson & Durieu 1855: 192) described the variey *Brachypodium distachyon* var. *platystachyon* from a name included in an exsiccatum collected by Balansa in 1852 (Plantes d’Algérie, 1852, No. 560). In the protologue was included a brief description, as “Var. γ. platystachyon. – Caulibus superne laevibus; spiculis latis, sub anthesi valde compressis, quam in planta vulgari majoribus; floribus longissime aristatis”, followed by the name “Syn. *Brachypodium distachyon* var. *platystachyon* Balansa *pl. Alger. exsicc.* n. 560” and the comment “Hab. In montosis, saepius promiscue cum planta vulgari, sed infrequens: *Oran*! *Saïda*! (Balansa)”.

We have found several syntypes preserved at B, MPU and GOET, from the gathering cited in the protologue as “*Brachypodium distachyon* var. *platystachyon* Balansa *pl. Alger. exsicc.* n. 560”. The herbarium sheets of these syntypes (i.e., B 10 0168260, GOET006090, MPU027897, MPU027896, MPU027895, W1889-0090643) bear an original printed label from the Balansa exsiccatum no. 560. This label is annotated as “B. Balansa, Pl. d’Algérie, 1852 / 560.

Brachypodium distachyon, R. S. / Var. platystachyon. / (B. Bal.) / Saïda, dans les terrains incultes” / 21 mai”. The lectotype of the name was designated by Scholz (Greuter & Raus 1998: 173) indicating the Balansa’s exsicata and the herbarium B. In B we only found one sheet B 10 0168260, which should therefore be considered the lectotype, while the other duplicated specimens would be isolectotypes.

According to our results, the names *Brachypodium hybridum* and *Trachynia platystachya* (≡ *Brachypodium distachyon* var. *platystachyon*) could be considered conspecific. Therefore, *Trachynia platystachya* is a new heterotypic synonyms of *B. hybridum*. However, *T. platystachya* predates *B. hybridum*, and the first name has priority and must be used when it and *B. hybridum* are considered synonyms. Nevertheless, the name *T. platystachya* is not known in the taxonomic or floristic literature. Therefore, its resurrection would create a problem in the nomenclature of this group of plants.

In conclusion, according to our studies, the names *Festuca rigida, Bromus pentastachyos, Brachypodium macrostachyum*, *Brachypodium megastachyum* and *Trachynia platystachya* ([≡ *Brachypodium distachyon* var. *platystachyon*) could be considered conspecific and attributable to the same taxon as *B. hybridum*. However, because *B. hybridum* has no priority over the other names under Art. 11.4 of the *ICN*, it is incorrect to consider these names as synonyms of *B. hybridum*.

Therefore, a proposal to conserve the name in current use *B. hybridum* against the disused or unused names *Festuca rigida*, *Trachynia pentastachya*, *Brachypodium macrostachyum*, *Brachypodium megastachyum* and *Trachynia platystachya* is needed.

***Brachypodium hybridum*** Catalán, Joch. Müll., Hasterok & Jenkins (2012: 402) **[*nom. cons. prop.*]**.

**Holotype**: Portugal, Lisboa, ABR113 inbred line, from seeds cultivated at Aberystwyth University, 30 May 2011, *Tim Langdon s.n.* (MA barcode MA 833766). **Isotypes**: JE barcodes JE00013253, JE00013254, K barcode K000975669, JACA code JACA-R298983).

= *Festuca rigida* Roth (1797: 12) **[nom. rej. prop.]**

**Lectotype** (**designated here**): s.l., s.d., *A.W. Roth s.n.,* B barcode B-W 02109-02 0 (Figure 5), probable isolectotypes: B barcodes B-W 02109-01 0 and B-W 02109-03 0.

**FIGURE 5.**
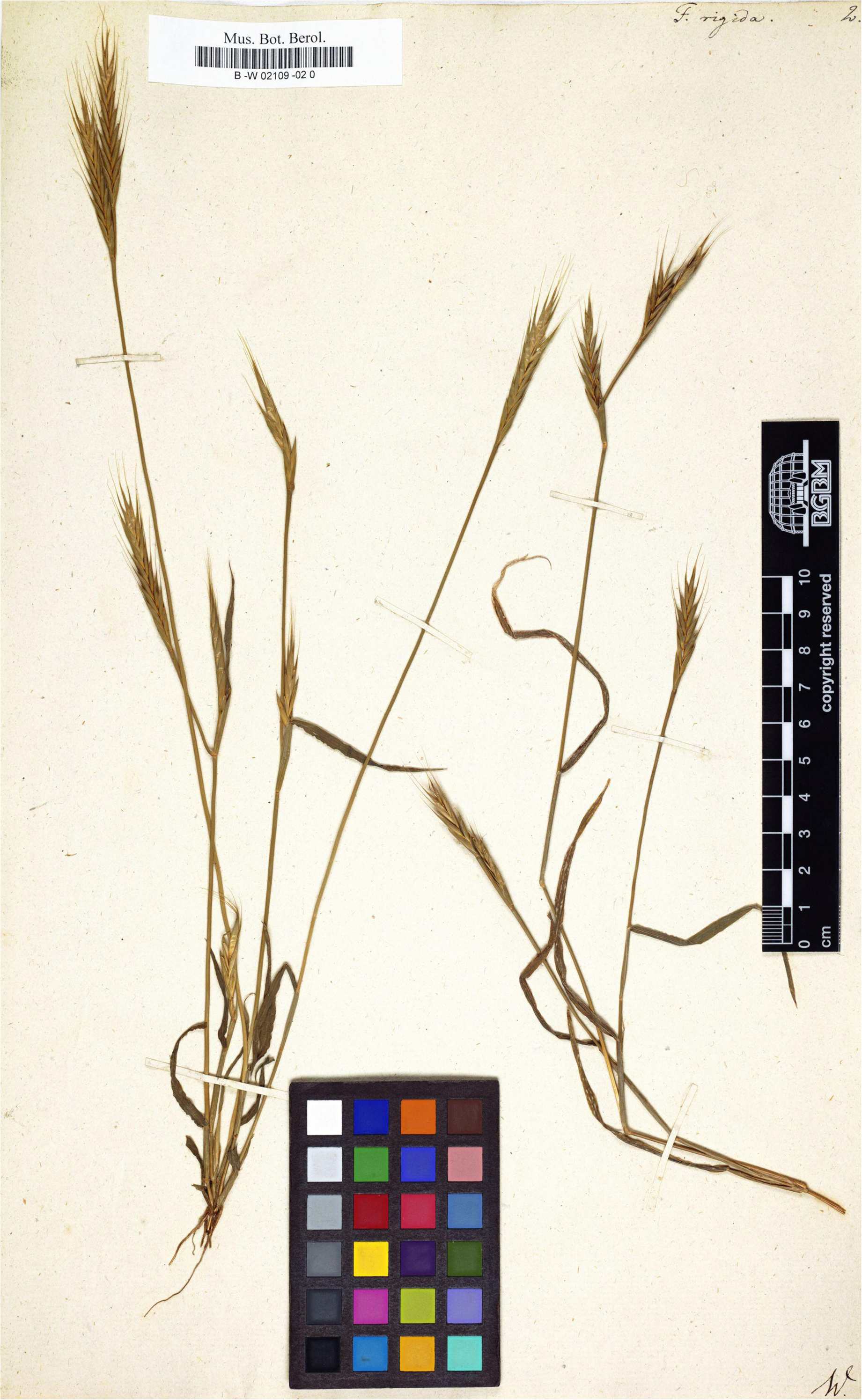
Lectotype of *Festuca rigida* Roth, B barcode B-W 02109-02 0. Image by courtesy of the herbarium B, reproduced with permission.

= *Trachynia pentastachya* (Tineo) Tzvelev (1975: 81) [≡ *Bromus pentastachyos* Tineo (1817: 4)]

[nom. rej. prop.]

**Lectotype** (designated by Steinberg in Bot. Jalrrb. Syst. 102: 411-425. 1981): PAL no. 19110 (Figure 6).

**FIGURE 6.**
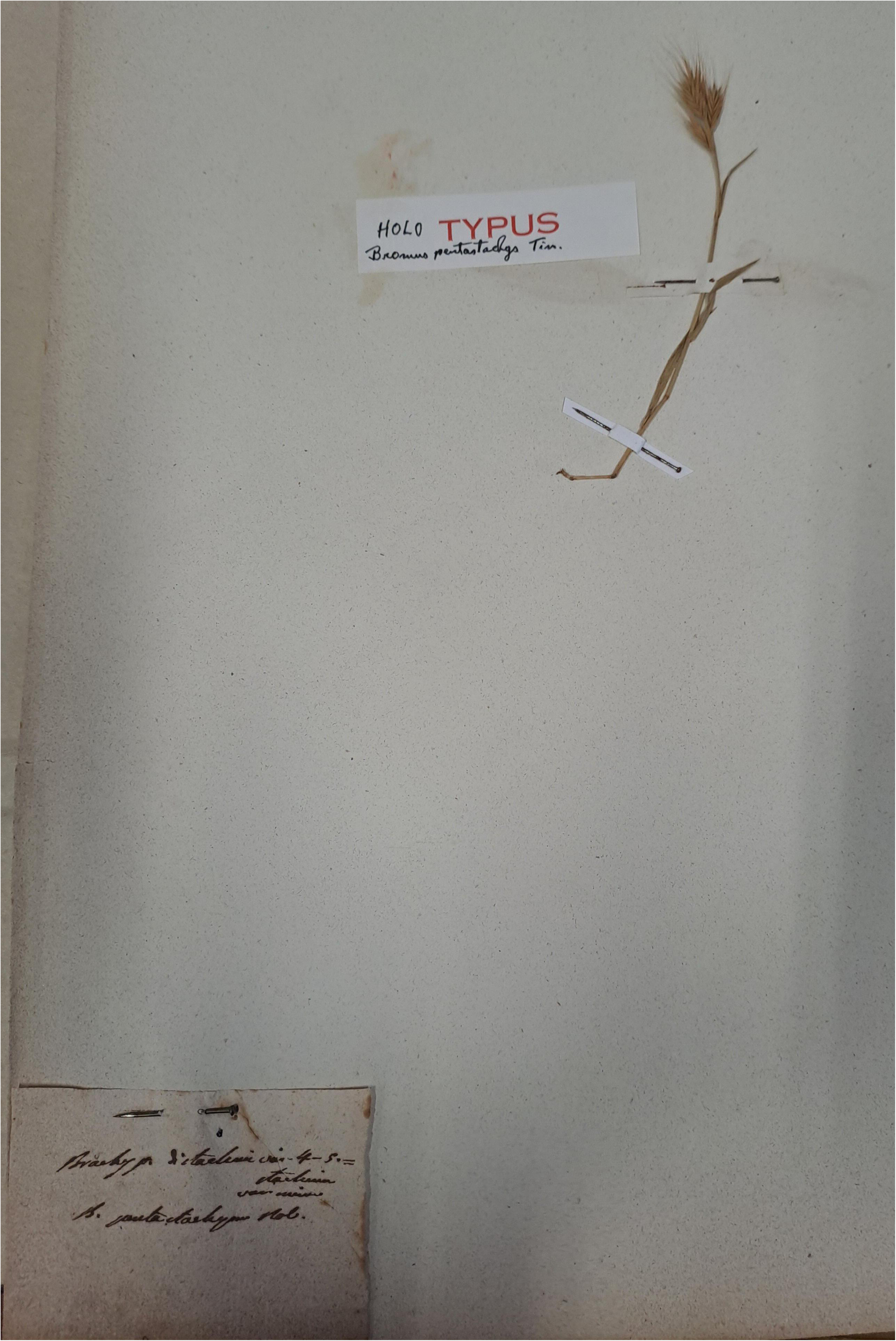
Lectotype of *Bromus pentastachyos* Tineo, PAL no. 19110. Image by courtesy of the herbarium PAL, reproduced with permission.

= *Brachypodium macrostachyum* Besser (1827: 651) **[nom. rej. prop.]**

**Lectotype** (designated by Schippmann in Boissiera 45: 178. 1991): [România, Bobâlna, Cluj], “Cult. Cremen. [Cremenea]”, s.d., *W. Besser s.n*., K barcode K000913822.

= *Brachypodium megastachyum* Besser (1827: 651) **[nom. rej. prop.]**

**Lectotype** (designated by Schippmann in Boissiera 45: 178. 1991): [România, Bobâlna, Cluj], “Cult. Cremen. [Cremenea]”, s.d., *W. Besser s.n*., K barcode K000913824.

Isolectotypes: LE barcodes LE00009972 and LE00009971.

= *Trachynia platystachya* (Balansa ex Coss.) H. Scholz in Greuter & Raus (1998: 173) [≡ *Brachypodium distachyon* var. *platystachyon* Balansa ex Coss. in Cosson & Durieu (1855: 192) [basionym]; ≡ *Brachypodium distachyon* subvar. *platystachyon* (Balansa ex Coss.) St.- Yves (1934: 481)] **[*nom. rej. prop.*]**

**Lectotype** (designated by Scholz in Greuter & Raus (1998:173)): [Algeria] “Saida dans les terrains incultes”, 215.1852, *Balansa, Pl. Algérie 1852, No. 560* (B [now barcoded B 10 0168260]). **Isolectotypes**: GH02433843, GOET006090, K003355093, L1223021, MPU027897, MPU027896, MPU027895, W1889-0090643.

***Brachypodium stacei*** Catalán, Joch.Müll., Hasterok & G. Jenkins (2012: 402 [18 of 21]

Another species of the *B. distachyon* complex is the diploid taxon *Brachypodium stacei.* This species is native to the circum-Mediterranean region (Greece, Israel, Italy (Sicily), Morocco, Palestine, Spain (including Balearic Islands: Formentera, Mallorca, and Menorca, and Canary Islands: Gomera and Lanzarote), and Tunisia (Catalán *et al*. 2012, López-Álvarez *et al*. 2015, Mu *et al*. 2023a, Campos *et al*. 2024). Its potential introduction into other regions where the species is non-native has yet to be confirmed. The name *B*. *stacei* is applied currently to an erect annual plant up to 44–76(115) cm high, leaf blades of culms (1.6)7–8(15.1) cm × (0.26)2–3(7) mm, erect to patent, flat, soft, curled, densely villose abaxially, with short and long hairs 0.1-0.7 mm, with panicle with (1)3–4(–5) spikelets, spikelets (13)22(41) cm, and lemmas (6.1)8–9(12.6) mm, with awn (7.5)12–13(18.2) mm (see Catalán *et al*. 2012, 2016a, 2016b).

The name *B. stacei* has also consistently been accepted and widely used in the scientific literature since its publication in 2012 (Shiposha *et al*. 2016, Sancho *et al*. 2018, 2022, Gordon *et al*. 2020, Díaz-Pérez *et al*. 2018, Gandour 2021, Scarlett *et al*. 2023, Mu *et al*. 2023a, Campos *et al*. 2024, Chen *et al*. 2024). In addition, NILs of the type specimen of *B. stacei* have been used to generate the reference genome of this species (*B. stacei* accession line ABR114; see Gordon *et al*. 2020).

***Triticum schimperi*** Hochst. ex Richard (1850: 441)

The protologue of *Triticum schimperi* Richard (in Tent. Fl. Abyss. 2: 441. 1850) includes the name “TRITICUM SCHIMPERI. Hochst., in *pl*. *Schimp*. *Abyss*., sect. I, n° 59”, a complete description in Latin, followed by the provenance “Crescit ad margines agrorum editiorum montis *Selleuda*, meridiem versus, mense Octobre (Schimper)”, and a comment “Observation. – C’est tout à fait le port du *Triticum ciliatum* DC. ou *Festuca distachya*. Mais les feuilles et le chaume sont cotonneux et blanchâtres, et les epillets ne m’ont paru conteir que de six à huit fleurs”.

The type of *T. schimperi* was indicated by Cufodontis (1968): “Typus: Schimper 59 (M. Selleuda, = Scholoda pr. Adua)”. Cufodontis’s indication could certainly satisfies the Art. 7.10 and 7.11 of the *ICN* (see Turland *et al*. 2018) and constitutes an effective lectotype designation, because he clearly indicated the “type element” mentioned in the Art. 7.11 (“… if the type element is clearly indicated…”) since an element can be considered as “…a single specimen or gathering…or illustration…” as indicated in the Art. 40.3 of the *ICN*. Moreover, the lectotypification proposed by Cufodontis could also be further narrowed to a single specimen by a “second-step” lectotypification according to Art. 9.17 of the *ICN* (Turland *et al*. 2018).

We have located several specimens that are part of the gathering mentioned in the protologue (in *pl*. *Schimp*. *Abyss*., sect. I, n° 59) (e.g., B 10 0168955, B 10 0168956, BM000922840, BR0000008367020, HAL0107050, HOH008866, K000367338, L1351188, LE00010059, LG0000090036248, MPU028011, P03328270, P03218973, S12-12906, S12-12907, TUB009106, W0030724, W0026339, W1889-0251135, WAG0001483.).

As the collection date is October, most of the material lacks complete inflorescences, or only contains a few spikelets. Only in two specimens (K000367338 and TUB009106) we have found complete material to be able to carry out the measurements. Among these specimens, we designate here as the lectotype of the name *Triticum schimperi* the specimen more complete and consequently more informative, preserved at TUB009106. The sheet TUB009106 bears five plants, with leaves and flowers, and a separate envelope with spikelets. The sheet also contains three labels, two handwritten label and a printed label from the exisccatum n° 59 of Schimper, this label is annotated as: “Schimperi iter Abyssinicum. / Sectio prima: plantae Adoënses. / 59. Triticum (Brachypodium) / Schimperi Hochst. / Ad margines agrorum editiorum montis Scholoda meridiem versus / U. i. 1840. / d. 3. Oct. 1837.”. The rest of the specimen of that area which are part of the exsiccatum n° 59 are isolectotypes.

*Triticum schimperi* has priority over *B. stacei* under Art. 11.4 of the *ICN.*, under the name *Brachypodium schimperi* (Hochst.) Chiov. (1919, n. s.xxvi. 83), and it would be incorrect to treat *B. schimperi* as the synonym of *B*. *stacei*. Therefore, a proposal to conserve the name in current use *B. stacei* against the disused or unused name *T. schimperi* is needed, or, altrenatively, a proposal to reject the name *T. schimperi* under Art. 56 of the *ICN*.

***Brachypodium stacei*** Catalán, Joch. Müll., Hasterok & Jenkins (2012: 402) **[*nom. cons. prop.*] Holotype**: Spain, Balearic Islands, Formentera, Torrent, ABR114 inbred line, from seeds cultivated at Aberystwyth University, 30 Oct 2010, *Luis Mur* (MA barcode MA833765, isotypes: JE barcode JE00013256, K barcode K000975666, JACA code JACA-R298982).

= *Triticum schimperi* Hochst. ex Richard (1850: 441) **[*nom. rej. prop.*]**

**Lectotype** ([first-step] designated by Cufodontis in Bull. Jard. Bot. Nat. Belg. 38(4): 1216. 1968): “Schimper 59 (M. Selleuda, = Scholoda pr. Adua)”; (second-step **designated here**): Ethiopia, Tigray, “Ad margines agrorum editiorum montis Scholoda meridiem versus U, i. 1840”, 3 Oct. 1837, *G.H.W. Schimper (Schimperi iter Abyssinicum. Sectio prima: plantae Adoënses. n. 59) W. Besser s.n*., TUB barcode TUB009106 (Figure 7). **Isolectotypes**: B 10 0168955, B 10 0168956, BM000922840, BR0000008367020, HAL0107050, HOH008866, K000367338, L1351187, L1351188, LE00010059, LG0000090036248, MHNF-HAFRIC36320, MICH1229144, MPU028011, P03328270, P03218973, S12-12906, S12-12907, W0030724, W0026339, W1889-0251135, WAG0001483.

**FIGURE 7.**
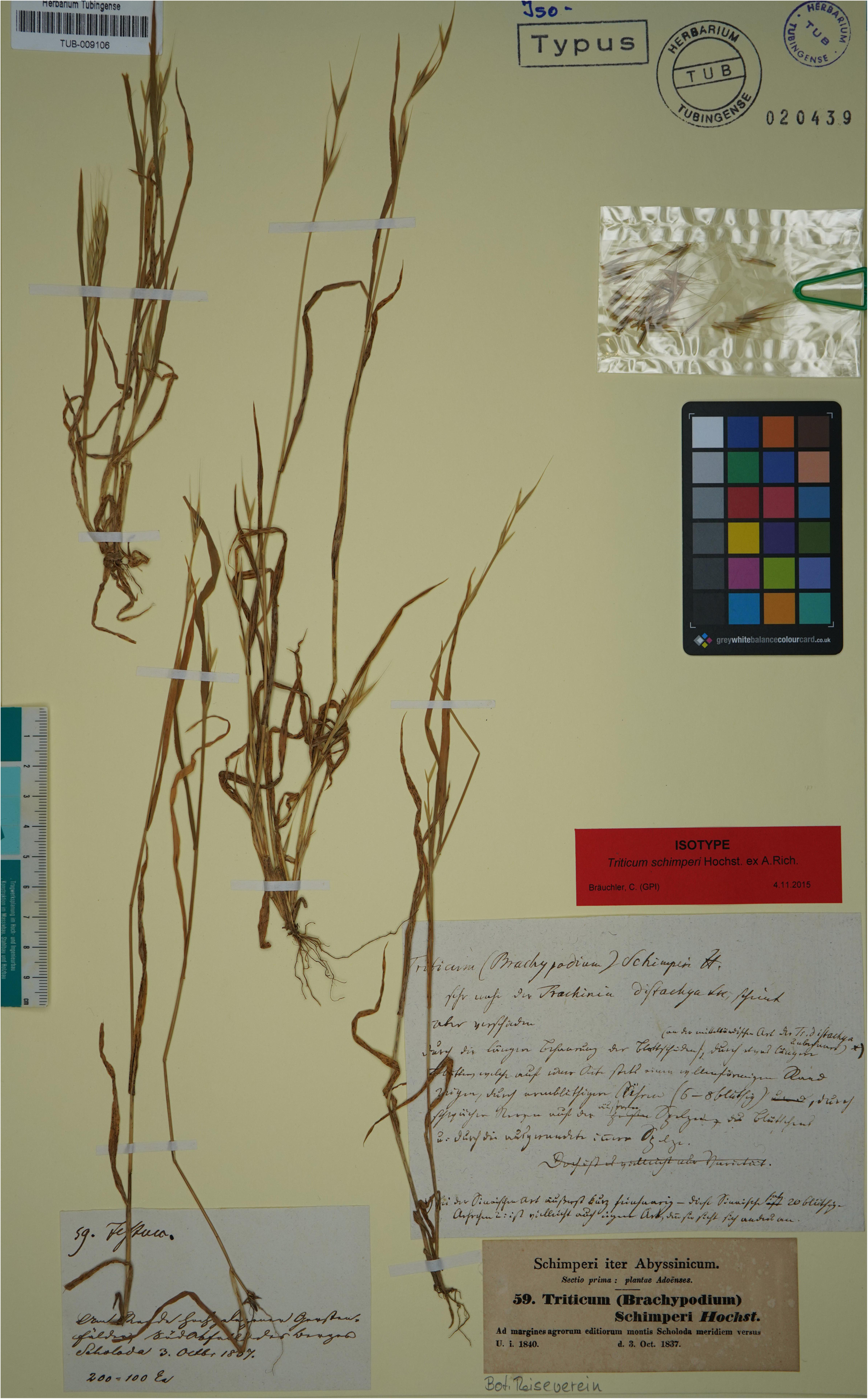
Lectotype of *Triticum schimperi* Hochst. ex Rich., TUB barcode TUB009106. Image by courtesy of the herbarium TUB, reproduced with permission.

### Other nomenclatural considerations in *Brachypodium distachyon* group

Eight names synonymized with *Brachypodium distachyon* sensu lato that lack original material are considered invalid or ambiguous names, or should be synonymized to other perennial *Brachypodium* species.

***Festuca ciliata*** Gouan (1762: 48), *nom. illeg*.

Although Schipmann (1991) lists it among the valid synonyms of *B. distachyon*, A. Gouan in his Hortus Regius Monspeliensis (Gouan 1762) mentions *Bromus distachyos* L. as a synonym.

Therefore, according to Art. 52.1 of the *ICN*, it is considered a superfluous name and is not included in the present study.

***Bromus pauper*** Schrank (1789: 370), *nomen ambiguum*

This is another name listed by Schipmann (1991) in its synonymy of *B. distachyon*. There is some controversy surrounding this name due to the asterisk preceding the proposal. This issue has been extensively studied by Slageren & Thomas Payne (2013), who, as previously noted by Schipmann (1991), argue that Schrank’s proposal (Schrank 1789) should be considered at a specific level.

Therefore, it can be regarded as a heterotypic synonym of *B. distachyon* sensu lato. In the description, Schrank mentions that he did not observe this species in Bavaria (hence the asterisk) and that he obtained it from a collection belonging to ‘Dr. Panzer’, presumably the German entomologist and botanist Georg Wolfgang Franz Panzer (1755-1829). We have been unable to find any herbarium material to study the characteristics of the proposal. Additionally, the plate by Christian Ehrenfried Weigel (1748-1831) in *Observ. Bot.: tab. I fig. 8* (1772), referring to his *Bromus distachyos*, depicts an inflorescence that does not help clarify which taxon it may represent. For all these reasons, and in the interest of maintaining the stability of the group, we consider this an obscure and ambiguous name (nomen ambiguum) and propose its rejection.

**FIGURE 8.**
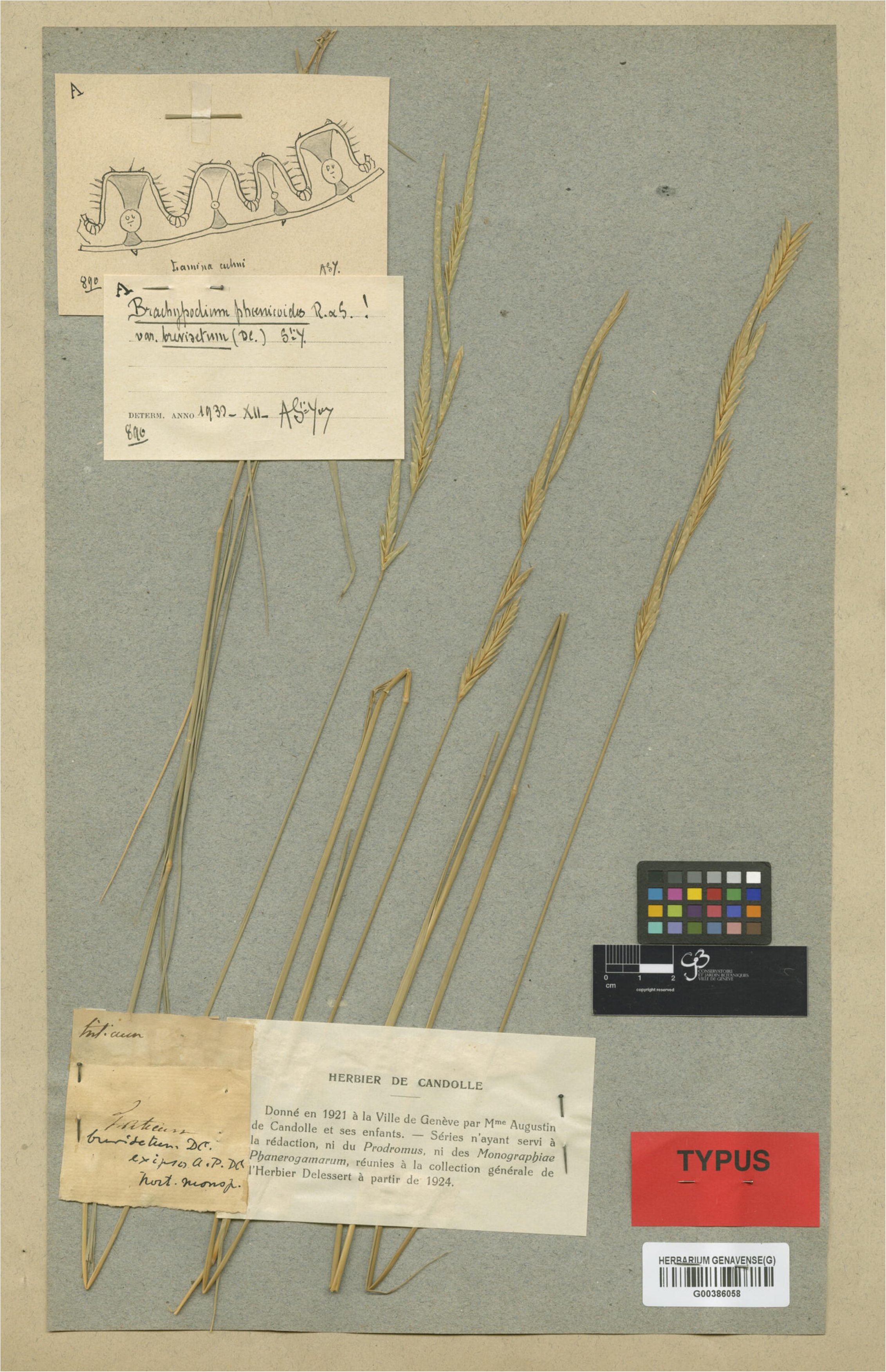
Lectotype of *Triticum brevisetum* DC., G barcode G00386058 [sheet 1] (the lectotype is mounted on two sheets). Image by courtesy of the herbarium G, reproduced with permission.

***Festuca pseudistachya*** Koeler (1802: 270), *nomen ambiguum*

Georg Ludwig Koeler, in his work *Descriptio Graminum in Gallia et Germania tam sponte nascentium quam humana industria copiosius provenientium* (Koeler 1802), includes a brief diagnosis, followed by the reference ‘Wigg. obs. bot. p. 16. t. I. f. 8’, and a complete description, concluding with the statement ‘Habit. in Bavar. Wirtemb. Pom. Austr.’. We have been unable to locate any original material that would allow us to study this name in relation to the annual species of *Brachypodium*. As with previous cases, the application of this name appears uncertain. It is a forgotten, ambiguous, and obscure name, which we propose for rejection as a nomen ambiguum in order to preserve the stability of the group.

***Triticum brevisetum*** Candolle (1813: 153)

Augustin Pyramus de Candolle publishes this name in his catalogue of the plants in the botanical garden of Monpellier (Candolle 1813), with a short description [“(221) Triticum brevisetum. T. spiculis 5-6 alternis teretiusculis 20-30-floris, flosculis breviter aristato-mucronatis, culmo ad apice ad basin scabro, foliis glaucis subconvolutis”], an information of unknown provenance (as, “Patria ignota”), and some indications that would make this species independent of *Triticum ciliatum* (Lam.) DC. [“Affine T. ciliato fl. fr. sed distinctum culmo ab apicè ad basin scabro nec loevi, spiculis 20-30-floris, nec 6-12-floris, aristis brevissimis, foliis subinvolucratis etc.”. In the Geneva herbarium at G there are two relevant gatherings that are part of the original material used by Candolle to describe his species. These gatherings are barcoded as G00386057 and G00386058 [with two sheets]. Schippman (1991: 177) mentioned for this name: “Patria incognita”, typum non vidi”. The sheet G00386057 bears a specimen with a leaf and inflorescences (image of the sheet available at https://www.ville-ge.ch/musinfo/bd/cjb/chg/adetail.php?lang=en&id=342994). The sheet bears a label with the name and locality: “*Triticum brevisetum* DC. / hort. monsp.” handwritten by Candolle. The gathering barcoded G00386058 is mounted in two sheets, the specimen is a complete material, with leaves, stems and inflorescences; the sheet also bears a label annotated as “Triticum brevisetum DC. / hort. monsp.”. The two specimens can be identified as belonging to a perennial species *Brachypodium* taxon, specifically to *B. retusum* Palisot de Beauvois (1812: 101). We therefore consider that this name should not be synonymized with the group of annual *Brachypodium* species. In conclusion, we designate as the lectotype of the name *Triticum brevisetum* the specimen barcoded as G00386058.

***Brachypodium retusum*** Palisot de Beauvois (1812: 101)

***=*** *Triticum brevisetum* Candolle (1813: 153) [**syn. nov.]**

**Lectotype** (**designated here**): [France]: Jardi des plantes de Montpellier, “Hortus Monspeliensis”", s.a., s.d., (G barcode G00386058 [two sheets]) (Figure 8). Probable isolectotype: G00386057.

***Triticum asperum*** Candolle (1813: 153), *nom. illeg*.

Another name publishes by Augustin Pyramus de Candolle in his catalogue of the plants in the botanical garden of Monpellier (Candolle 1813) is *Triticum asperum* Candolle (1813: 153), which we have encountered in several of the databases consulted, where it is listed as a valid heterotypic synonym of *B. distachyon*. A review of the prologue reveals that A.P. de Candolle compares his species to *Festuca rigida* Roth, making this name superfluous under Art. 52.1 of the ICN.

***Bromus paradoxus*** Presl (1820: 23), *nomen ambiguum*

Karel Bořivoj Presl publishes the name *Bromus paradoxus* Presl (1820: 23) accompanied with a brief description, indicating its origin: ‘Hab. In arenis fluminis magni (Fiume grande)’ and the comment ‘Peculiare forsan genus constituit’. Despite extensive efforts, no source material related to C. Presl’s proposal has been found, either in the Prague herbaria containing specimens of this author (PR, BRNM) or in other herbaria where some of his collections may be held. Given the limited information in the brief description, it is difficult to associate this name with any specific species. On the other hand, given the absence of original material and the fact the name has not been treated in over two centuries, its application is clearly uncertain. It is recommended to be rejected as a nomen ambiguum.

***Triticum flabellatum*** Tausch (1837: 117), *nomen ambiguum*

Ignaz Friedrich Tausch describes this taxon with a brief description, but without indicating its origin or referring to any other name or illustration. We contacted the curators of the PRC Herbarium, where most of Tausch’s collection material is held (Stafleu & Cowan 1983), but nothing was found, neither for this name nor for other new proposals by Tausch within the genus *Triticum*. One possible explanation is that the plants were cultivated in the garden of Joseph von Canal, who did not produce any herbarium sheets (Patrik Mráz, pers. obs.). Further consultation with other herbaria that may hold Tausch’s material has also been unsuccessful. Therefore, given the absence of original material its application is clearly uncertain. It is recommended to be rejected as a nomen ambiguum.

***Brachypodium geniculatum*** Koch (1848: 422), *nomen ambiguum*

Another of the names for which we have not found original material to study and compare, and thus assign to one of the described species of annual *Brachypodium* taxa is *B. geniculatum*, a species described on the basis of material from the province of Yerevan in Armenia, collected by Koch himself, and material from Sicily, collected by A. Dietrich and studied by Koch in the Berlin herbarium. Much of K. Koch’s herbarium was destroyed in a fire of B herbarium in 1943 (Lack 1978), and probably these collections disappeared. However, since this is an obscure, unknown and unused name, we also believe that it should be proposed for rejection as a nomen ambiguum.

### Final conclusions

According to our study (see Table 2), the names Festuca rigida, Bromus pentastachyos, Brachypodium macrostachyum, Brachypodium megastachyum and Trachynia platystachya would have priority over B. hybridum. On the other hand, the current name B. stacei is threatened by the name Triticum schimperi.

**TABLE 2.**
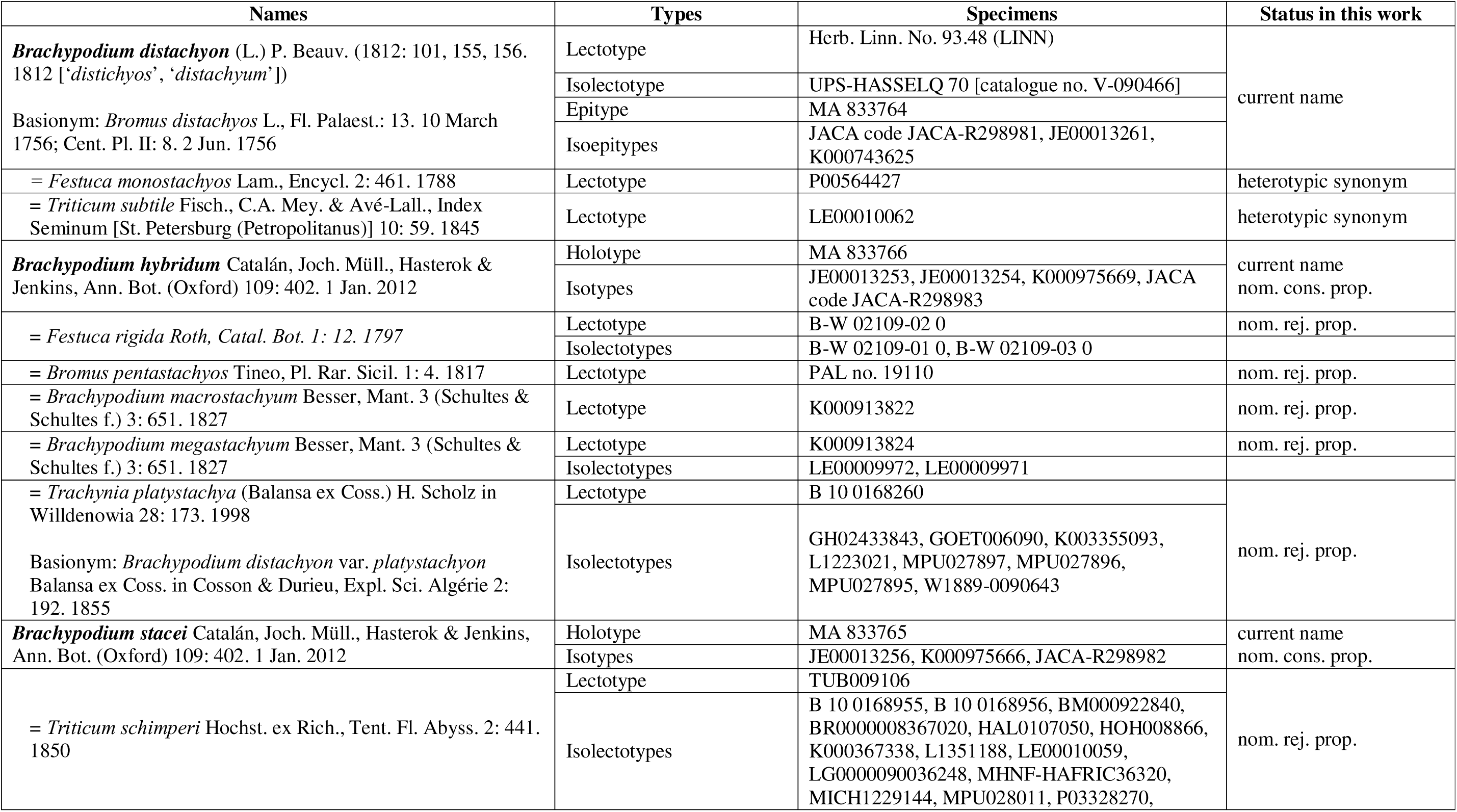
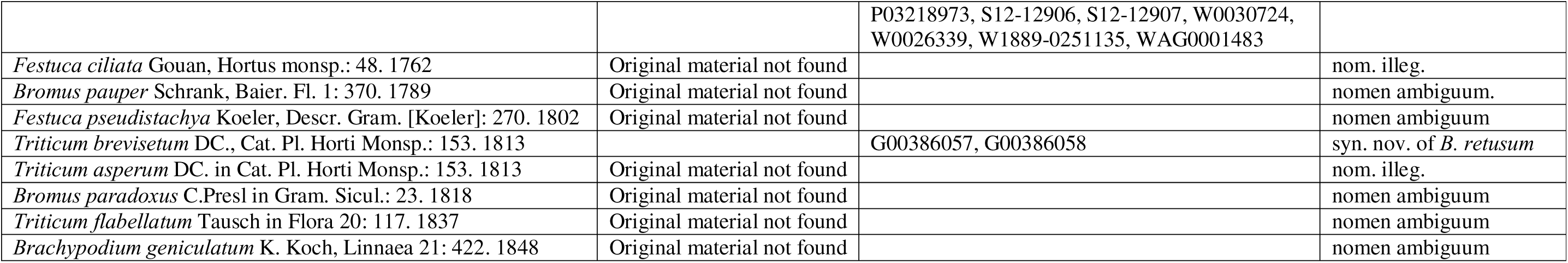
Taxonomical and nomenclatural synthesis, with the names associated to the *Brachypodium distachyon* (L.) P. Beauv. complex, types and taxonomical attribution according to the present study.

Maintaining the priority rules of the International Code of Nomenclature for algae, fungi, and plants (Shenzhen Code) within the B. distachyon complex group (including B. hybridum and B. stacei) will lead to name and status changes. This is undesirable for these model species systems commonly referenced in the most recent literature as B. hybridum or B. stacei. Much confusion will be created if the name B. hybridum is replaced by Festuca rigida, Brachypodium macrostachyum, or Brachypodium megastachyum, which is undesirable and represents a problem for the nomenclatural stability of B. hybridum. The same will occur if the name B. stacei is replaced by Triticum schimperi.

Therefore, in our parallel publications we submit the respective proposals of conservation for B. hybridum and B. stacei, the most stable and commonly used names for the group of names presented above. The present work serves as a solid base to uncover these nomenclatural proposals. Furthermore, the study of seven names traditionally synonymized with B. distachyon has proven uninformative due to the lack of original material for study. As a result, these names remain obscure and confusing, and we believe a proposal for their rejection would be beneficial. Two of these names (i.e., Festuca ciliata and Triticum asperum) are considered illegitimate, while five others (i.e., Bromus pauper, Brachypodium geniculatum, Festuca pseudistachya, Bromus paradoxus, Triticum flabellatum), due to the lack of original material and the uncertain application of these names, are regarded as nomina ambigua.

Finally, one name synonymized with B. distachyon, Triticum brevisetum, has been found to represent species more closely related to other perennial Brachypodium species. We propose it should be synonymized with B. retusum.

## Acknowledgements

The authors would like to thank the B, BM, BR, G, GH, GOET, HAL, HOH, JACA, JE, K, L, LE, LG, LINN, MA, MHNF, MICH, MPU, P, PAL, PRC, S, TUB, UPS, and W herbaria for providing the images of the studied types, Patrik Mráz for his help in the search for specimens of Tausch, Gianniantonio Domina for consulting bibliographic works difficult to access, and Miguel Campos for the R script. Thanks are also due to Dr. John Wiersema and Dr. John McNeill for their advice, assistance, and valuable comments. This research was supported by the Spanish Ministry of Science and Innovation (Grants No. TED2021-131073B-I00, PDC2022-133712-I00, and PID2022-140074NB-I00), and the Spanish Aragon Government-European Social Fund Bioflora (Grant No. A01-23R) to PC.

## Availability of data and materials

Herbarium materials of the non-type specimens of *Brachypodium distachyon*, *B. stacei* and *B. hybridum* used in this study are available at the University of Zaragoza (UZ) herbarium. Documents containing the input and output files for the SPSS discriminant analysis and the custom R script designed for the two-dimensional plotting of the DA function values are available on Github (https://github.com/Bioflora/Brachy_nomenclature).

---

**TABLE S1.**
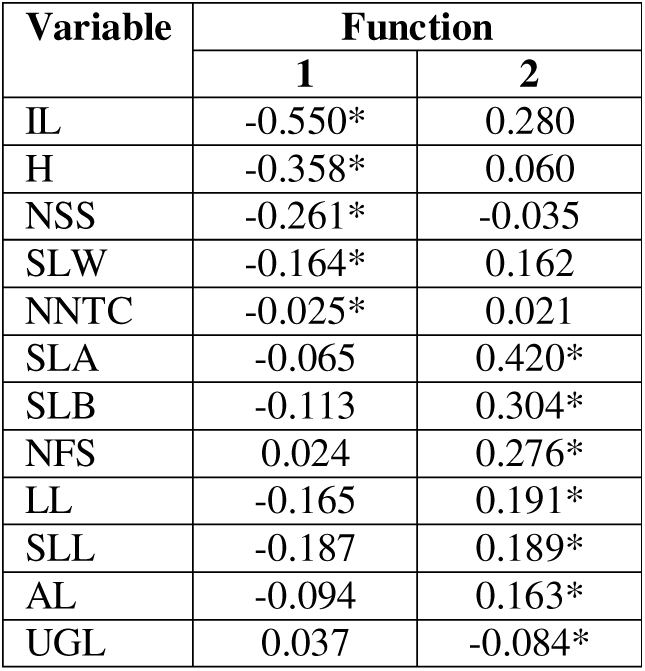
Contribution of morphological variables to the first canonical discriminant functions obtained for the data set of 50 individuals of *B. distachyon*. *B. stacei* and *B. hybridum* and 12 specimen types analyzed. Functions 1 and 2 accumulated 71.5% and 28.5% of variance, respectively. The Variables that contributed more to the functions are highlighted in bold. Abbreviation of variables: H (Plant height). NNTC (Number of nodes of tallest culm). SLL (Second leaf length from the base of the plant). SLW (Second leaf width). IL (Inflorescence length). NSI (Number of spikelets per inflorescence). SLA (Spikelet total length. without awns). SLB (Spikelet length from the base to the apex of the fourth lemma. without awns). NFS (Number of flowers per inflorescence). UGL (Upper glume length). LL (Lemma length from the basal floret). AL (Awn length. the longest within the spikelet).

